# PKM2-mediated epigenetic reprogramming regulates hypoxic expression of *PFKFB3* to promote breast cancer progression

**DOI:** 10.1101/2022.11.06.515384

**Authors:** Madhura R. Pandkar, Adarsh Raveendran, Kajal Biswas, Srinivas Abhishek Mutnuru, Jharna Mishra, Atul Samaiya, Shyam K. Sharan, Sanjeev Shukla

**Affiliations:** Department of Biological Sciences, Indian Institute of Science Education and Research Bhopal, Bhopal, Madhya Pradesh 462066, India; Center for Cancer Research, National Cancer Institute, Frederick, MD 21702-120; Department of Pathology, Bansal Hospital (BH), Bhopal, Madhya Pradesh 462016, India; Department of Surgical Oncology, BH, Bhopal, Madhya Pradesh 462016, India

**Keywords:** Breast cancer, hypoxia, epigenetics, glycolysis, PKM2

## Abstract

The hypoxic milieu is a critical modulator of aerobic glycolysis, yet the regulatory mechanisms existing between the key glycolytic enzymes in hypoxic cancer cells are largely unexplored. In particular, M2 isoform of pyruvate kinase (PKM2) − the rate-limiting enzyme of glycolysis, is well-known to confer adaptive advantages under hypoxia. Herein, we report a non-canonical role of PKM2 in functioning as a co-activator of HIF-1α to govern the transcription of 6-phosphofructo-2-kinase/fructose-2,6-biphosphatase 3 (*PFKFB3*). Nuclear PKM2 enhances HIF-1α and p300 occupancy at *PFKFB3* hypoxia responsive elements (HREs) resulting in its upregulation. Consequently, absence of nuclear PKM2 fails to recruit HIF-1α which activates an opportunistic occupancy of HIF-2α at *PFKFB3* HREs. Enhanced binding of HDAC3 also occurs in the absence of PKM2 which prevents HIF-2α from efficiently inducing PFKFB3 to hamper proliferation of hypoxic breast cancer cells. In addition, clinical relevance of the study has been investigated by demonstrating that Shikonin blocks nuclear translocation of PKM2 to suppress *PFKFB3* expression. Furthermore, MCF7 cells-derived xenograft tumors in mice exhibited substantial tumor growth inhibition when treated with shikonin, highlighting the vitality of targeting PKM2. Taken in concert, this work provides novel insights into contributions of PKM2 in modulating hypoxic transcriptome and a previously unreported molecular axis exhibited by the hypoxic breast cancer cells for ensuring the maintenance of PFKFB3 expression essential for achieving high glycolytic flux.

**Statement of significance:** Nuclear PKM2 orchestrates the binding of histone modifiers to epigenetically alter PFKFB3 promoter and affects the binding of HIF-2α. Notably, targeting this axis attenuates proliferation of hypoxic breast cancer cells.

## Introduction

Glycolysis, the central pathway of cellular metabolism, is highly responsive to various environmental cues. Numerous reports published in the last few decades have revealed that cancer cells are extensively dependent on glycolysis even under optimal oxygen concentrations, a phenomenon referred to as the Warburg effect (1, 2). Moreover, frequently observed tumor-associated microenvironmental condition of hypoxia has been well-studied to prompt the genesis of drastic metabolic alterations, further enhancing the glycolytic potential of cancer cells (3, 4). The adaptative strategies exhibited by hypoxic cancer cells are primarily manifested by the activity of a family of transcription factors known as hypoxia inducible factors (HIFs). The HIFs are heterodimeric proteins comprising of an oxygen-sensitive α subunit and an oxygen insensitive β subunit. Under optimal O_2_ concentration, the HIF-1α subunit undergoes prolyl hydroxylation (by prolyl hydroxylases, i.e., PHDs) at the residues Pro 402 and 564 (5), targeting it for ubiquitin-dependent proteasomal degradation (6). However, under hypoxia, PHDs are functionally less active, due to which the prolyl hydroxylation of HIF-1α is substantially blocked. Consequently, the stabilized transcription factor translocates to the nucleus and pairs with its constitutive partner, HIF-1β. This functional heterodimer drives the expression of a plethora of genes by binding to the consensus sequence (5’RCGTG3’) known as hypoxia responsive element (HRE) to govern various cellular pathways (7–9). One of the most striking effects of hypoxia can be seen on the metabolism of cancer cells. In this context, glycolysis has extensively been investigated to undergo alterations under hypoxia, as numerous glycolytic genes are HIF1 targets (4).

Interestingly, PKM2 exhibits incredible capacity to perform various non-canonical functions dictated by diverse microenvironmental cues (10) and is also investigated to enhance cancer cell survival under hypoxia (11). One of the most imperative non-canonical feature of PKM2 is its ability to translocate into the nucleus to act as a transcriptional co-activator of HIF-1α (12). Subsequently, it has been demonstrated that the moonlighting activities associated with PKM2 have pivotal role in shaping the metabolic landscape of hypoxic cancer cells. However, role of PKM2 in modulating the hypoxic transcriptome remained elusive due to the lack of appropriate model systems that are selectively knockout for PKM2 and not PKM1 in the context of cancer.

As the BORIS-binding site (BBS) present in exon 10 of pyruvate kinase (*PKM*) gene has a dictatorial influence in promoting biased expression of PKM2 over PKM1 (13), we mutated the BBS to generate PKM2 knockout breast cancer cell lines. Therefore, if for a gene PKM2 acts as a transcriptional activator, its hypoxic induction will be affected in the BBS mutants (BBS Mut) as compared to their wild-type BBS (WT BBS) counterpart cells. HIF-1α ChIP-seq paired with transcriptome array analysis revealed *PFKFB3* as a previously unidentified target of the PKM2-HIF-1α axis. Moreover, PKM2 was observed to promote enhanced binding of p300 to cause transcriptional upregulation of *PFKFB3*. Interestingly, the cells lacking HIF-1α or PKM2 expression did not exhibit a complete loss of PFKFB3, and HIF-2α was identified as a non-canonical regulator, which maintains only basal-level expression in the absence of HIF-1α or PKM2. We also observed enhanced occupancy of HDAC3 in absence of PKM2, which prevented HIF-2α from inducing PFKFB3 expression. Notably, this adaptive switch was found to significantly enhance the dependency on OXPHOS, ultimately compromising the proliferative potential of hypoxic breast cancer cells.

We also systematically delineate that Shikonin− a pharmacological inhibitor of PKM2, interacts with R399/400 residues to block its nuclear translocation, thus leading to the disruption of PKM2-HIF-1α-PFKFB3 axis. Moreover, our *in-vivo* findings clearly demonstrate that shikonin hampers the growth of breast cancer cells expressing PKM2 compared to their PKM2 knockout counterpart cells. We envisage that targeting this axis has therapeutic potential to inhibit proliferation of hypoxic breast cancer cells by substantial diminution of glycolytic rate.

## Materials and Methods

### 1. Cell culture

Human breast cancer cell lines MCF7 and HCC1806 were obtained from American Type Culture Collection (ATCC). MCF7 and HEK293T were cultured at 37ºC, 5% CO2 in DMEM (Invitrogen, 12800017, lot no. 2248833). HCC1806 was cultured in RPMI-1640 (Invitrogen, 23400021, lot no. 2144859). The culture media for all the cell lines was supplemented with 10% fetal bovine serum (FBS; Sigma, F7524, lot. no BCBX8466), 100 units/ml of penicillin and streptomycin (Invitrogen, 15140122, lot no. 2321120), and 2mM/l L-glutamine (Invitrogen, 25030081, lot no. 1917006). The BBS Mut MCF7 cell line was generated as previously mentioned (13). The media and cell culture conditions for various CRISPR/Cas9 mutants were kept similar as the wild-type cells. Cell lines were subjected to hypoxia treatment of 1% oxygen in a Ruskinn INVIVO2 400 hypoxia chamber.

### 2. Plasmids

The mCherry-tagged PKM2 overexpression construct was generated as previously mentioned (13). The PKM2 overexpression lentiviral plasmid was constructed by cloning the PKM2 insert between the NotI and EcoRI sites of the plasmid pSCALPS_ZSgreen kindly gifted by A. Chande (IISER Bhopal). The primers used for generating the insert are mentioned in Supplementary table 2.

### 3. Quantitative RT-PCR

Total RNA was extracted using TRIZol reagent (Ambion, 15596018, lot no. 260712) as per the manufacturer’s instructions. The concentration was determined using Eppendorf BioSpectrometer, and 1μg of total RNA was used to synthesize cDNA using SuperScript® III First-Strand Synthesis System (Invitrogen, 18080-051, lot no. 2291381). Amplifications were performed in duplicates using GO taq QPCR master mix (Promega, A6002, lot no. 0000385100) and light cycler 480 II (Roche) according to the manufacturer’s protocol. The primers were designed using the IDT PrimerQuest tool (https://www.idtdna.com) and are enlisted in Supplementary table 1. The average cycle thresholds from three independent biological replicate samples were calculated and normalized to housekeeping control gene *RPS16* using the formula 2^ ^(Ct_control − Ct_target)^. Student’s t-test was used to compare gene/exon expression between two different groups. *P*< 0.05 was considered statistically significant. The primers used in qRT-PCR analysis are mentioned in Supplementary table 2.

### 4. Immunoblotting

The cells were lysed using urea lysis buffer (8M urea, 2M thiourea, 2% CHAPS, 1% DTT) supplemented with 1× protease inhibitor cocktail (PIC; leupeptin 10–100 μM, pepstatin 1μM, EDTA 1–10 mM, AEBSF <1 mM) at 4°C for 30min and spun at maximum speed (16,900×g) in a 4°C centrifuge for 2hrs. The supernatant was separated, quantified and an equal concentration of protein samples was loaded for every experiment. The well-separated proteins were electro-transferred on an activated PVDF membrane. After transfer, the blots were incubated with recommended dilutions of primary antibodies overnight at 4°C, followed by 1hr incubation with secondary antibody. The blots were scanned using Odyssey membrane Scanning system. Quantification of the bands was performed using GelQuant software (version 1.8.2). The technical details of the antibodies used are enlisted in Supplementary table 3.

### 5. PFK assay

WT BBS and BBS Mut cells subjected to normoxic or hypoxic treatment were lysed using assay buffer provided in the Phosphofructokinase (PFK) Activity Colorimetric Assay Kit (Sigma, MAK093, lot no. 6E31K07760). An equal quantity of protein lysates was used to determine the total PFK activity as per the manufacturer’s instructions. The readings were measured using a microplate reader (BioTek Eon, 11-120-611) set at 37°C and an optical density of 450nm. The values are represented as mean ± SD of duplicates from a representative experiment. The statistical significance was calculated using unpaired t-test.

### 6. Lactate Assay

Post-treatment, equal number of WT BBS and BBS Mut MCF7 and HCC1806 cells were lysed using ice-cold lactate assay buffer provided in the Lactate assay kit (Sigma, MAK064-1, lot no. 3F09K06270). The lactate quantification was performed using the deproteinized lysates by following the manufacturer’s instructions. The readings were measured at room temperature using a microplate reader set at an optical density of 450nm. The statistical significance was calculated using unpaired t-test.

### 7. α-Ketoglutarate assay

WT and BBS Mut cells of MCF7 and HCC1806 cells subjected to 24hrs of hypoxia were processed for α-ketoglutarate estimation. Briefly, equal number of cells were lysed using assay buffer provided in the alpha KG assay kit (abcam, ab83431, lot no. GR3410863-1). The lysates were then deproteinized using perchloric acid and nutralized using 2M KOH as per manufacturer’s instructions. Finally, an equal amount of sample was taken and the reaction was incubated at 37°C for 30min. The absorbance was measured using a microplate reader set at an optical density of 450nm. The statistical significance was calculated using unpaired t-test.

### 8. Chromatin immunoprecipitation (ChIP) assay

ChIP assay was performed as previously described (14). Briefly, about 10 million cells were crosslinked and scraped in PBS, followed by lysis and sonication. About 25μg of sheared chromatin (approx length 250-500bp) was immunoprecipitated with an antibody of interest following overnight incubation at 4°C. Immunoprecipitated protein–DNA complexes and 5% input were analyzed by qRT-PCR using GO taq QPCR master mix (Promega, A6002, lot no. 0000385100) in triplicate using primers specific for *PFKFB3* HREs. Additionally, each experiment was performed at least thrice. Normalization was performed using the formula [2^^ (Ct input – Ct immunoprecipitation)^]. The obtained values were further normalized to the relative rabbit IgG and control IP values. Significance between the two groups was calculated using student’s t-test, with a value of <0.05 being considered statistically significant. HIF-1α ChIP-seq data was analyzed as previously described (15). Over-representation analyses (ORA) for GO Biological processes were generated using ShinyGo 0.76.2 (16). False discovery rate (FDR)-value of <0.05 was considered significant while obtaining enriched GO terms. Top 20 enriched terms were represented using dot plot and ranked according to the fold enrichment value. A heat map was constructed using the online tool Morpheus (https://software.broadinstitute.org/morpheus) for the 44 common genes between HIF-1α ChIP-seq and the DEGs obtained from BBS Mut vs WT BBS Hypoxia condition.

### 9. RNA interference

The cell lines used throughout this study were plated at a seeding density of 3×10^5^ cells per well of a six-well culture plate and allowed to attach for 24hrs. The lentivirus containing small hairpin RNA (shRNA) (Sigma, Mission Human Genome shRNA Library) against the target gene was inoculated in the presence of 8μg/ml polybrene (Sigma, H9268, lot no. SLBH5907V) containing media. Cells were selected for 72hrs using 1μg/ml puromycin (Sigma, P9620, lot no. 034M4008V) and subsequently used for various experiments. The sequence of shRNAs used in this study is provided in Supplementary table 1.

### 10. Human Transcriptome Array 2.0

Total RNA was isolated from WT BBS and BBS mutant MCF7 cells using TRIZol (Ambion, 15596018, lot no. 260712) reagent and PureLink RNA Mini Kit (Invitrogen, 12183025, lot no. 1862249) as per manufacturer’s instructions. The concentration was determined using Eppendorf BioSpectrometer, and 100ng of total RNA was used for biotinylated cDNA synthesis using GeneChip™ WT Plus Reagent Kit (Invitrogen, 902281, lot no. 01059768) as per manufacturers protocol. Following amplification, cDNA fragmentation was performed, and 5.5μg of fragmented cDNA was hybridized on Affymetrix GeneChip™ Human Transcriptome Array 2.0 (HTA2.0) chips (Invitrogen, 902162, lot no. 4364589) for 16 hrs at 45°C. The chips were washed and stained using the Affymetrix Fluidics Station 450. Post hybridization, the fluorescence intensity of the arrays was scanned using the Affymetrix Scanner 7G. The raw files generated in CEL format after scan were further used for analysis.

### 11. Human Transcriptome Array 2.0 data analysis

The CEL files were analyzed using Transcriptome Array Console 4.0 (Invitrogen, version 4.0.2.15) using the gene+exon–SST-RMA method of summarization. Genes with thresholds of absolute fold-change >2, *P*<0.05, and false discovery rates (FDRs) <0.05 were selected as differentially expressed genes (DEGs). Venn diagram was generated using Venn Diagram Plotter software developed by Pacific Northwest National Laboratory (https://omics.pnl.gov/software/venn-diagram-plotter). The volcano plot for DEGs was plotted using GraphPad Prism 9 software.

### 12. Immunofluorescence

Cells were plated at a seeding density of 3×10^4^ on a coverslip in a 12-well plate. After treatment, the cells were washed thrice with ice-cold PBS followed by 4% formaldehyde fixation and subsequent permeabilization with 0.1% Triton X-100 solution for 15min. Post blocking for 1hr at RT using 2% BSA solution, cells were incubated with appropriate primary antibodies overnight at 4°C. Cells were then washed thrice with PBS and incubated with Alexa-Flour 555 anti-rabbit IgG secondary antibody for 1hr at RT. Subsequently, the cells were counterstained with DAPI (Invitrogen, D1306, lot no. 1673432) and mounted using fluoroshield (Sigma, F6182, lot no. MKCN2676). For Ki67 staining, primary antibody incubation was directly followed by nuclei counterstaining using Hoechst 33342 (Invitrogen, HI399, lot no. 1932847). Imaging was performed using Olympus FV3000 confocal laser scanning microscope with a 60× Plan Apo N objective (oil, 1.42 NA), and image analysis was performed using Image J software.

### 13. Breast cancer sample collection

Prior to utilizing tumor specimens for research purposes, informed consent was obtained from all the patients. Paraffin-embedded tumor and adjacent normal breast tissue sections affixed on poly-L-lysine-coated slides were collected from Bansal Hospital, Bhopal, India. The study was approved by the Institute Ethics Committee of the Indian Institute of Science Education and Research Bhopal, India.

### 14. Immunohistochemistry

Breast tumor tissues of 5 microns size were obtained from Bansal hospital, Bhopal for immunohistochemical analysis. Briefly, the sections were deparaffinized at 65°C followed by xylene treatment and rehydration using ethanol gradients. Antigen retrieval was then performed by boiling the slides at 98°C in citrate buffer (pH 6.0) for 10min. Endogenous peroxidase was quenched using hydrogen peroxide solution for 15min at RT followed by blocking using 5% BSA for 1 hour at RT. Primary antibodies against the proteins of interest were incubated overnight at 4°C. Subsequently, secondary antibody detection and DAB staining were performed using Biogenex super sensitive™ polymer HRP IHC detection kit (QD430-XAKE, lot no. QD4300919) as per manufacturer’s instructions. The details of patient samples are enlisted in Supplementary table 4.

### 15. Subcellular fractionation

Subcellular fractionation was performed as described previously (17). Briefly, cells were scraped and collected in PBS followed by centrifugation at 3000rpm for 5min. The pelleted cells were resuspended in cytoplasmic lysis buffer (10mM HEPES [pH 8.0], 1.5mM MgCl_2_, 10mM KCl, 0.5% NP-40) supplemented with 1× PIC and incubated on ice for 15min. The resultant mixture was pelleted by centrifugation at 3000rpm, 4°C for 10min. The supernatant was collected as cytoplasmic fraction and stored at -80°C until further use. The pellet was washed with PBS followed by resuspension in nuclear lysis buffer (10mM HEPES [pH 8.0], 1.5mM MgCl_2_, 400mM NaCl, 0.1mM EDTA, 20% Glycerol) supplemented with 1× PIC. The mixture was vortexed vigorously and was incubated at 4^°^C for 30min. The mixture was then centrifuged at maximum speed (16,900 × g) for 30min at 4^°^C. The supernatant was collected as nuclear fraction.

### 16. Establishment of CRISPR/Cas9 knockout cells

The CRISPR/Cas9-mediated HIF-1α knockout of HCC1806 was generated as previously mentioned (14). For generating HIF-2α knockouts, exon 2 of *EPAS1* (gene coding for HIF-2α) targeting sgRNA was designed using GPP sgRNA Designer tool (https://portals.broadinstitute.org/gpp/public/analysis-tools/sgrna-design). The sgRNA was cloned using the BbsI site of lentiviral vector pLentiCRISPR-E (Addgene, 78852) as described (https://media.addgene.org/cms/filer_public/6d/d8/6dd83407-3b07-47db-8adb-4fada30bde8a/zhang-lab-general-cloning-protocol-target-sequencing_1.pdf). The sequence of the obtained construct was verified using Sanger sequencing. Lentivirus containing sgRNA cloned plasmid was generated in HEK293T. Post transduction, cells were subjected to puromycin (1μg/mL) selection for 5-6 days. Subsequently, single-cell suspension was performed using a serial-dilution method and seeded in a 96-well plate. The positive knockout clones were identified using immunoblotting and the sequencing analysis of the region targeted by sgRNA. A similar protocol was followed for generating BBS Mut HCC1806 cells. The sequence of the sgRNAs used is provided in Supplementary table 1.

### 17. Co-Immunoprecipitation

The hypoxia-treated MCF7 and HCC1806 cells were lysed using cell lysis buffer (10mM Tris [pH 7.5], 150mM NaCl, 0.5mM EDTA, 0.5% NP-40) supplemented with 1×PIC for 30min. The lysates were then centrifuged at the highest speed for 20min. The supernatants were collected, quantified, and incubated with anti-PKM2, anti-HIF-1α, and Normal Rabbit IgG antibodies overnight at 4°C. 25μL Protein G Dynabeads™ (Invitrogen, 10004D, lot no. 01013573) were washed thrice with cell lysis buffer, added to the immunoprecipitated lysate and further incubated for 2hrs at 4°C. The immunoprecipitated complex was then isolated by magnetizing the beads and washed four times with cell lysis buffer. Post washing, the beads were boiled with 2× Laemmli buffer for 5min. The beads were then separated, and the eluted proteins were analyzed with immunoblotting.

### 18. Luciferase reporter assay

103bp, 89bp, and 75bp fragments spanning *PFKFB3* HRE1, HRE2, and HRE3 respectively were individually amplified and cloned between KpnI and XhoI sites of the pGL3 basic expression vector (Promega, E1751) using MCF7 genomic DNA as a template. The primers used to amplify the fragments are enlisted in supplementary table 2. Cells were seeded in a 24-well plate and allowed to attach. The wells were co-transfected with different *PFKFB3*-HRE luciferase constructs along with pRL-TK renilla luciferase plasmid (Promega, E2231). 12hrs post-transfection, the cells were subjected to appropriate treatments for 12hrs. Subsequently, the cells were lysed, and the luciferase activity was determined using GloMax-Multi Detection System (Promega) by normalizing to the Renilla luciferase activities.

### 19. Cell Proliferation assay

The cell lines were seeded at a density of 85×10^4^ cells per well of a 12-well plate. Post completion of time points, the dye exclusion method of viable cell counting using trypan blue dye (Himedia, TC193, lot no. 0000190691) and hemocytometer was employed.

### 20. MitoTracker™ Deep Red staining

MitoTracker™ Deep Red (Invitrogen, M22426, lot no. 2112250) stock solution was prepared as per the manufacturer’s instructions. 1×10^5^ cells were seeded in a well of 12-well plate and allowed to attach for 24hrs. 70nM working solution of MitoTracker™ Deep Red made using phenol red-free DMEM (Gibco, 21063-029, lot no. 1967601) or phenol red-free RPMI (Gibco, 11835-030, lot no. 1951096) supplemented with 10% FBS was added to the cells. After 45min of incubation in the dark at 37°C, the cells were washed with PBS and stained for nucleus using Hoechst 33342. Subsequently, the cells were trypsinized, washed twice with PBS, finally resuspended in PBS, and subjected to flow cytometry using fluorescence-activated cell sorting (FACS) Aria III by Becton Dickinson. Analysis of the data was performed using FlowJo software (version 10.7.1). For fluorescence imaging, MitoTracker™ Deep Red was added to the 3D-spheroid culture at a final concentration of 150nM followed by incubation in the dark at 37°C. The staining was then visualized using Olympus FV3000 confocal laser scanning microscope with a 60× Plan Apo N objective (oil, 1.42 NA).

### 21. Mitochondria isolation from cell lines

Mitochondria were isolated from the breast cancer cells as mentioned previously (18). Briefly, cells were washed once with ice-cold PBS. Subsequently, the cells were scraped in PBS and centrifuged at 600g at 4°C for 10min. After discarding the supernatant, the pellet was resuspended in 2-3 mL of ice-cold incubation buffer ([Ibc] 10mM Tris–MOPS, 10mM EGTA/Tris, 200mM sucrose [pH-7.4]). The cells in resuspension were partitioned in multiple microcentrifuge tubes for homogenization using a hand-operated micro pestle, and the resultant homogenate was spun at 600g for 10min at 4°C. The supernatant was collected and centrifuged at 7000g for 10min at 4°C. The obtained mitochondrial pellet was washed with 200μL of ice-cold Ibc and finally resuspended in the minimal volume of the buffer that remained after discarding the supernatant.

### 22. Mitochondrial Complex I assay

The isolated mitochondrial lysate was quantified, and an equal amount of the preparation was used to estimate the mitochondrial complex I activity as per the manufacturer’s protocol (Sigma, MAK359, lot no. 6E01K09680). The readings were measured at an optical density of 600nm using a microplate reader set at RT. The values are represented as mean ± SD of technical duplicates from a representative experiment. The statistical significance was calculated using unpaired t-test.

### 23. IC_50_ calculations

3×10^3^ cells were seeded in each well of a 96-well cell culture plate. The cells were allowed to attach for 24hrs. Appropriate serial dilutions of Shikonin (Cayman chemicals, 14751, lot no. 0566727-12) stocks were made in cell culture media and added to the wells. 24hrs post-treatment, a 10μL aliquot of 2mg/mL tetrazolium dye, 3-(4,5-dimethylthiazol-2-yl)-2,5-diphenyltetrazolium bromide (Alfa Aesar, L11939, lot no. 10196058) dissolved in culture medium was added to each well and incubated for 2-3hrs at 37°C. The formazan crystals were then solubilized using 100μL DMSO, and the absorbance was measured at 570nm using a microplate reader. The IC_50_ value was determined using GraphPad Prism 9 software.

### 24. Molecular docking studies

Global docking analysis for shikonin with PKM2 was performed using AutoDock Vina (19). The protein structure of PKM2 (20) (PDB ID: 1T5A) was downloaded from the RCSB Protein Data Bank (21), cleaned, and processed in PyMOL 2.5 (22). The chemical structure of shikonin was downloaded from the PubChem (PubChem ID: 479503), and energy optimization was performed using Chem3D 20.1 software. The protein and ligand files were prepared using AutoDockTools (ADT). Polar hydrogens, Kollman charges, and water molecule deletion were assigned using ADT and saved in PDBQT format. The grid size was set to 86 × 80 × 87 along the axes with a spacing of 1.000A°, and the grid center was designated at X = 70.806, Y = -2.988, and Z = 90.970 to cover the entire protein surface. The docking parameters of exhaustiveness, number of modes, and energy range were set at 8, 20, and 4 kcal/mol, respectively. Rigid docking of PKM2 and shikonin was scored, and binding modes were generated using AutoDock Vina. The different binding modes of the docking were energy-optimized using the YASARA Energy Minimization server (23). The optimized poses were finally visualized and analyzed using PyMOL 2.5, Discovery Studio Visualizer v21.1.0, and Protein-Ligand Interaction profiler (PLIP) (24).

### 25. Generation of 3-dimensional spheroids

The 3D spheroids were generated using an overlay method as previously described (25, 26) with few modifications. 50μL of Growth Factor Reduced (GFR) Basement Membrane Matrix (Corning, 356230, lot no. 9343006) was spread uniformly in a well of 96-well cell culture plate and allowed to solidify at 37°C. 7×10^3^ cells were resuspended in 100μL of appropriate cell-culture media containing 1μg/mL hydrocortisone (Sigma, H0888, lot no. SLBG4963V) and 5μg/mL insulin (Sigma, I1882, lot no. SLBR1114V) and added over the solidified matrix. Media containing 10% GFR matrigel was then subsequently overlayed as the topmost layer and incubated at 37°C in the presence of 5% CO_2_. Hormones containing media was replaced every 72hrs.

### 26. Hypoxia detection in 3D spheroids

Hypoxic regions in the 3D spheroids were detected using Hypoxyprobe™-1 (HP-1000, lot no. 111318) as described previously (27). Briefly, pimonidazole hydrochloride was added to the cell culture media at a final concentration of 100μM and incubated at 37°C for 2hrs. The samples were fixed using 4% paraformaldehyde and subjected to immunofluorescence protocol. The samples were finally incubated with an anti-pimonidazole mouse IgG1 monoclonal antibody that binds to protein, peptide, and amino acid adducts of pimonidazole in hypoxic cells. The hypoxic regions were visualized by using Alexa-Flour 488 anti-mouse IgG.

### 27. Animal experiments

6×10^6^ cells (WT BBS and BBS Mut MCF7 cells) in 100 μl of PBS was injected s.c. into the flanks of 6-week-old female athymic nude mice. One week prior to injection of cells, estradiol cypionate (Depo-Estradiol Covetrus NA, 074962) in cottonseed oil was administered s.c. at 1mg/kg between the shoulder blades using 25G needle. Estradiol injections was given once every week. Once the tumors reach a size of 100 mm^3^, mice were randomly assigned to four groups. The resource equation method was used to calculate the size of group and no selection criteria was used to assign groups. Shikonin (Sigma, 074962) dissolved in DMSO or DMSO only were injected in mice i.p either 1.5mg/kg of Shikonin in 20μl DMSO or 20μl DMSO every other day for 2 weeks. Tumor volume was measured every other day for 5 weeks. Tumors were harvested before they reach the maximal size (2000 mm^3^) permitted by NCI-Frederick ACUC. Tumor sizes were measured by digital Vernier Caliper and tumor volume (in mm^3^) was calculated using the formula (length×width^2^)/2. Tumor growth inhibition for day 31 was calculated using the formula TGI %= (1-[mean volume of treated tumors at day 31/ mean volume of control tumors at day 31]) ×100. Mice were maintained as per ACUC guidelines.

### 28. Statistical Analysis

All statistical analyses were performed using GraphPad Prism 7. Unless otherwise mentioned, all data are represented as mean ±SEM analyzed using two tailed Student’s t-test, unpaired t-test for the metabolic assays, and Welch’s test for the animal experiments. The statistical methods for each analysis are described within the figure legends or Materials and methods. P value less than 0.05 was considered significant. *P ≤ 0.05, **P ≤ 0.01, ***P ≤ 0.001, ****P ≤ 0.0001, ns= not significant.

### 29. Data availability

All the data supporting the findings are included in the article and its supporting information.

## Results

### 1. Hypoxia promotes nuclear translocation of PKM2 in breast cancer cells

Similar to other glycolytic enzymes, PKM2 performs various non-metabolic functions besides catalyzing the conversion of phosphoenolpyruvate (PEP) to pyruvate (28–31). One of the most intriguing non-canonical features of PKM2 is its ability to undergo nuclear translocation and functioning as a co-activator of transcription factors (12, 32). To verify if hypoxia serves as a stimulus for nuclear translocation of PKM2 in breast cancer cells, we subjected MCF7 and HCC1806 to 1% O_2_ for 24hrs. Immunostaining analysis visualized using confocal microscopy (Fig. S1.A, Fig. S1.B) and immunoblot assay (performed using nuclear and cytoplasmic protein lysates) revealed that post 24hrs of hypoxic treatment, there was a drastic increase in the nuclear pool of PKM2 (Fig. S1.C). To understand the molecular mechanism behind this phenomenon, we performed a lentivirus-mediated knockdown of Ran− a GTPase previously reported to aid nuclear migration of PKM2 under glucose restriction stress (32). The distribution of PKM2 upon exposing the Ran knockdown cells to 24hrs of hypoxia assessed using immunofluorescence imaging indicated a reduced pool of nuclear PKM2 compared to the control cells. Our results highlight that Ran plays a vital role as a nuclear translocator of PKM2 under hypoxic stress as well (Fig. S1.D, S1.E). Furthermore, to understand the role of PKM2 in orchestrating transcriptional alterations in hypoxic breast cancer cells, we further confirmed its interaction with HIF-1α by performing co-immunoprecipitation (Co-IP) assay. We found that endogenous PKM2 was specifically precipitated by the anti-HIF-1α antibody (Fig. S1.F) and consequently, upon performing a reverse pulldown using PKM2 antibody, HIF-1α was also observed to be precipitated (Fig. S1.G). Altogether, these results indicate that hypoxia serves as a stimulus for nuclear translocation of PKM2, which then interacts with HIF-1α to exert its non-canonical functions in breast cancer cells.

### 2. Nuclear PKM2 regulates the expression of metabolism-related genes under hypoxia

We have previously demonstrated the crucial involvement of epigenetically-acting transcription factor BORIS in dictating the alternative splicing of *PKM* (13). Upon introducing mutations in the BORIS-binding site (BBS) present in the exon 10 region of *PKM* using the CRISPR/Cas9, we generated PKM2 knockout cell lines of MCF7 and HCC1806, which expresses only PKM1 isoform (Fig. 1A, Fig. 1B).

**Figure 1.**
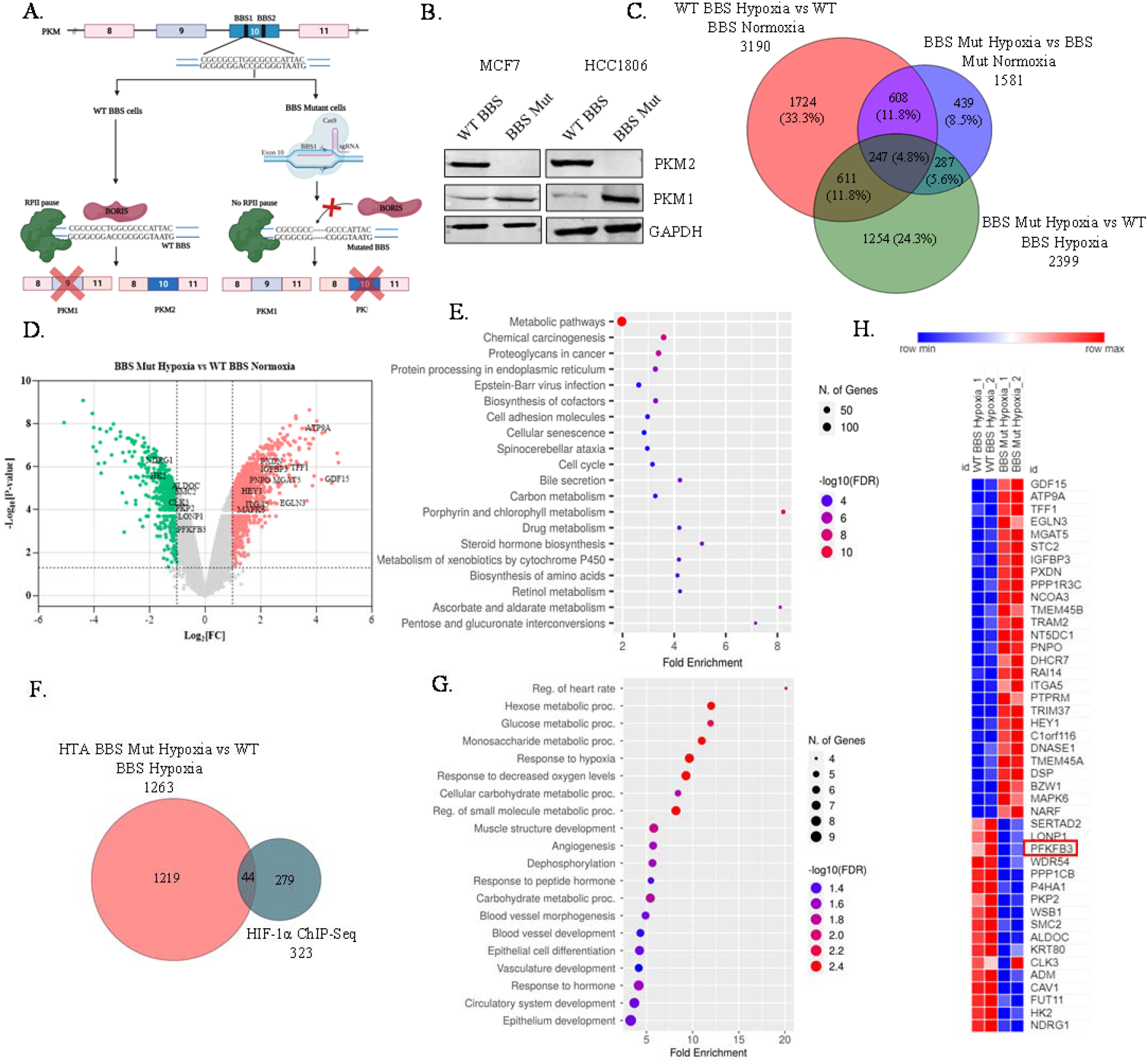
Nuclear PKM2 affects transcriptome of hypoxic breast cancer cells. (A) Schematic representation of strategy employed for generating CRISPR/Cas9-mediated PKM2 knockout cell lines. (B) Immunoblot analysis of PKM2 and PKM1 in WT BBS and BBS Mut cell lines of MCF7 and HCC1806. (C) Venn diagram depicting the DEGs under the conditions of WT BBS MCF7 normoxia vs. hypoxia, BBS Mut MCF7 normoxia vs. hypoxia, and BBS Mut MCF7 hypoxia vs. WT BBS MCF7 hypoxia. (D) Volcano plot of HTA 2.0 analysis showing the DEGs in BBS Mut MCF7 hypoxia vs. WT BBS MCF7 hypoxia. The significantly downregulated and upregulated genes are indicated with green and red points respectively (P < 0.05 and Fold Change >2). The top targets have been highlighted. (E) Dot plot representing top 20 enriched terms for biological processes for the DEGs obtained from BBS Mut Hypoxia vs WT BBS Hypoxia HTA 2.0 analysis. (F) Venn diagram depicting the common genes between HIF-1α ChIP-seq and BBS Mut hypoxia vs. WT BBS hypoxia HTA 2.0 analysis. A total of 44 HIF-1α target genes were differentially expressed due to absence of PKM2. (G) Dot plot representing the top 20 enriched terms for GO Biological Process (FDR < 0.05) in the common genes identified from HIF-1α ChIP-Seq and DEGs from BBS Mut Hypoxia vs WT BBS Hypoxia HTA 2.0. (H) Heat map representation of 44 common genes from HIF-1α ChIP-Seq and DEGs from HTA 2.0. For (B) representative images are provided. Error bars show mean values ± SD (n = 3 unless otherwise specified). As calculated using two-tailed Student’s t-test, *P ≤ 0.05, **P ≤ 0.01, ***P ≤ 0.001, ****P ≤ 0.0001.

Next, to examine the global transcriptomic alterations induced by PKM2 depletion, we performed Human Transcriptome Array 2.0 (HTA 2.0) analysis in wild-type BBS (WT BBS) and BBS mutant (BBS Mut) MCF7 cells subjected to 24hrs of hypoxia. Data analysis performed using Transcriptome Analysis Console (TAC) revealed that 3190 and 1581 genes were differentially expressed in WT BBS normoxia vs. WT BBS hypoxia (condition 1), and BBS Mut normoxia vs. BBS Mut hypoxia (condition 2) respectively. Moreover, 855 genes were observed to be common between condition 1 and condition 2 (Fig. 1C). Notably, 2399 genes were identified to be differentially expressed in BBS Mut hypoxia vs. WT BBS hypoxia (condition 3) from which 1430 genes were upregulated and 969 genes were downregulated. The significant events of differentially expressed genes (DEGs) in BBS Mut hypoxia vs. WT BBS hypoxia with Fold Change >2 and P < 0.05 are marked in Fig. 1D. The GO terms for biological processes enrichment analysis of DEGs in condition 3 revealed that metabolism-related genes were prominently enriched indicating that PKM2 has paramount influence in altering the expression of metabolism-related genes (Fig. 1E).

To further identify the hypoxia-specific genes that are regulated by PKM2, we analysed ChIP-seq data of HIF-1α in MCF7 and compared the target genes with the DEGs obtained from condition 3. 44 genes were observed to be common between the 323 HIF-1α target genes and the 1263 coding DEGs from condition 3 (Fig.1F). In addition, the GO terms for biological processes such as hexose metabolic process, glucose metabolic process, and response to hypoxia were distinctly enriched in the 44 common genes (Fig. 1G). A heat map for the HIF-1α targets affected by absence of PKM2 is presented in Fig. 1H.

Conclusively, our data indicates the vitality of the non-canonical nuclear function of PKM2 in promoting global transcriptomic alterations of the glucose metabolism-associated genes under hypoxia.

### 3. PKM2 is required for the hypoxic induction of *PFKFB3*

To validate the potential novel targets of the HIF-1α-PKM2 axis obtained from HTA 2.0 analysis, we performed a qRT-PCR screen of 15 glycolytic genes that are reported HIF-1α targets (Fig. S1.H). Conclusively, our results suggested that 6-phosphofructo-2-kinase/fructose-2,6-biphosphatases 3 (PFKFB3) requires PKM2 for its hypoxic induction. PFKFB3 catalyzes the reversible reaction of converting fructose 6-phosphate (Fru 6 P) to fructose 2,6-bisphosphate (Fru 2,6 BP) (33, 34), latter being potential regulator of phosphofructokinase-1 (PFK-1) activity. As PFK-1 catalyzes the committed step of glycolysis, its constitutive activation results in an elevated glycolytic rate (Fig. 2A). In addition to possessing the highest kinase: bisphosphatase activity compared to other family members (35), PFKFB3 has been positively associated with cancer progression and aggression (36–38).

**Figure 2.**
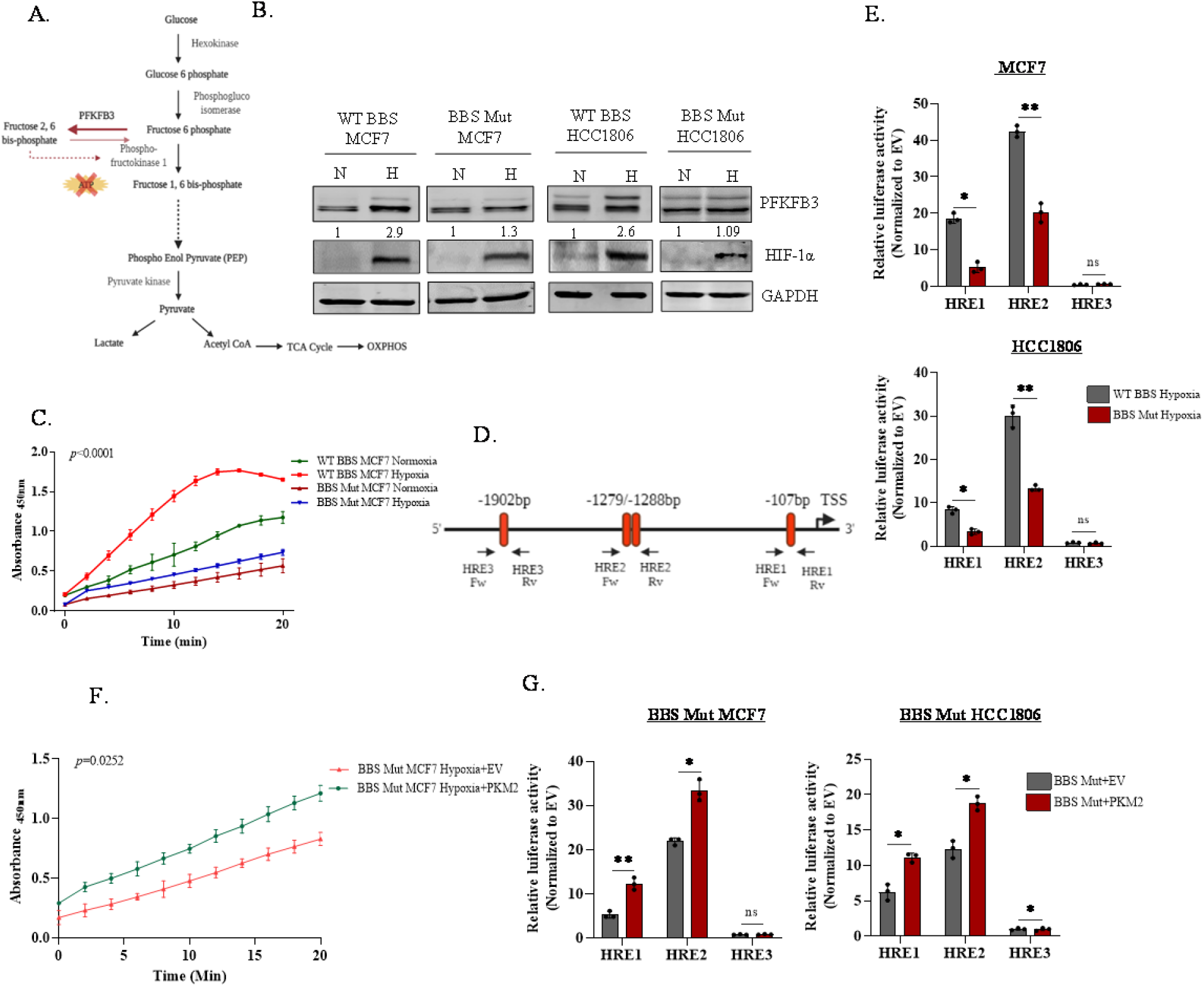
PKM2 regulates PFKFB3 expression under hypoxia. (A) Schematic representation of the role of PFKFB3 in glycolysis. PFKFB3 catalyzes the reversible reaction of converting fru 6 P to fru 2,6 BP to regulate the activity of PFK-1. (B) Immunoblot analysis depicting the hampered hypoxic induction of PFKFB3 in BBS Mut cell lines of MCF7 and HCC1806. (C) PFK assay performed in WT BBS and BBS Mut MCF7 exposed to normoxia or 24hrs hypoxia. (D) Schematic representation of the location of the three HREs in the *PFKFB3* promoter. The arrows indicate the primers used in ChIP-qRT PCR and for amplifying respective HREs for luciferase construct generation. (E) Luciferase assay performed in WT BBS and BBS Mut MCF7 and HCC1806 cells show the dampened activity of HRE1 and HRE2 in the mutant cells highlighting their functional involvement in *PFKFB3* regulation under hypoxia. (F) PFK assay results depicting enhanced PFK activity of the BBS Mut MCF7 cells upon re-introduction of PKM2 under hypoxia. (G) PKM2 overexpression followed by luciferase assay in BBS Mut cell lines of MCF7 and HCC1806 cells showing increased FLuc activity of HRE1 and HRE2 under hypoxia. For (B), (C), and (F), representative images are provided. For luciferase assays, error bars show mean values ± SD (n = 3 unless otherwise specified). As calculated using two-tailed Student’s t-test and unpaired t-test for PFK assays, *P ≤ 0.05, **P ≤ 0.01, ***P ≤ 0.001, ****P ≤ 0.0001.

Immunoblot analysis validated that hypoxic induction of PFKFB3 was hampered in the BBS Mut cell lines compared to their respective WT BBS cells (Fig. 2B). Consequently, to check if the BBS Mut MCF7 cell lines exhibited a decreased phosphofructokinase (PFK) activity as well, PFK assay was performed. The results clearly indicated that owing to a difference in PFKFB3 expression, the BBS Mut MCF7 cells manifested a lower PFK activity under hypoxia (Fig. 2C). As discussed previously, Ran aids nuclear translocation of PKM2; thereby, we observed a reduction in *PFKFB3* induction in both the WT BBS cell lines used in our study upon performing Ran knockdown under hypoxia (Fig. S2.A, S2.B).

To identify the regulatory region which drives PFKFB3 expression in PKM2-HIF-1α-dependent manner, we explored the functional contribution of each HREs located at the positions -107 (HRE1), -1279/-1288 (HRE2), and -1902 (HRE3) relative to the transcriptional start site (TSS) (39) (Fig. 2D). The three fragments of the length 103bp, 89bp, and 75bp spanning HRE1, HRE2, and HRE3 respectively were individually cloned upstream of a firefly luciferase (FLuc) coding sequence in a pGL3 basic vector. The reporter constructs were co-transfected with pRL-TK vector (containing renilla luciferase expression controlled by the SV-40 promoter) to the cell lines, followed by exposure to 1% O_2_ for 12hrs. The FLuc activity was significantly decreased in the BBS Mut cell lines of MCF7 and HCC1806. Conclusively, HRE1 and HRE2 were identified as the functional *PFKFB3* HREs from our luciferase assay data (Fig. 2E).

Notably, re-introduction of PKM2 in BBS Mut MCF7 cells lead to its hypoxia-specific nuclear translocation (Fig. S2.C) paired with a heightened induction of PFKFB3 (Fig. S2.D). Consequently, the rescue effect observed in PFKFB3 expression was also accompanied by rise in hypoxic PFK activity (Fig. S2.E, Fig. 2F). Luciferase assay also indicated a rescue effect in the HRE1 and HRE2 activity (Fig. 2G). Altogether, our data conclusively suggests that HRE1 and HRE2 are the critical sites that are regulated by nuclear PKM2.

### 4. PKM2 is an essential interacting partner of HIF-1α in transcriptionally regulating *PFKFB3*

To further verify luciferase assay results, ChIP assay was performed with HIF-1α and PKM2 antibody in the WT BBS and BBS Mut MCF7 cell lines exposed to 24hrs of 1% O_2_. As evident from the prominent enrichment of the aforementioned factors, we concluded that HRE1 and HRE2 are functionally active sites for PKM2-HIF-1α binding (Fig. 3A). Additionally, to investigate if HIF-1α is an obligate interacting partner of PKM2, we generated a stable HIF-1α knockout cell line of HCC1806 (H1AKO). PFKFB3 expression and PFK activity was observed to be dampened under hypoxia in H1AKO cells (Fig. S3.A-S3.D). Furthermore, ChIP assay corroborated a complete lack of PKM2 occupancy in H1AKO cells confirming that PKM2 itself is unable to bind *PFKFB3* HREs under hypoxia (Fig. 3B). Additionally, the FLuc activity of HRE1 and HRE2 was also drastically reduced in H1AKO cells (Fig. S3.E). To understand the mechanism underlying PFKFB3 induction by PKM2, we performed ChIP assay of histone acetyltransferase p300 which is primarily involved in transactivating HIF-1α target genes (40). Both p300 occupancy and the histone modification it catalyzes-H3 lysine-9 acetylation (H3K9Ac) was drastically reduced in BBS Mut cells showing that PKM2 induces epigenetic alteration, making *PFKFB3* HREs accessible for transcriptional initiation (Fig. 3C).

**Figure 3.**
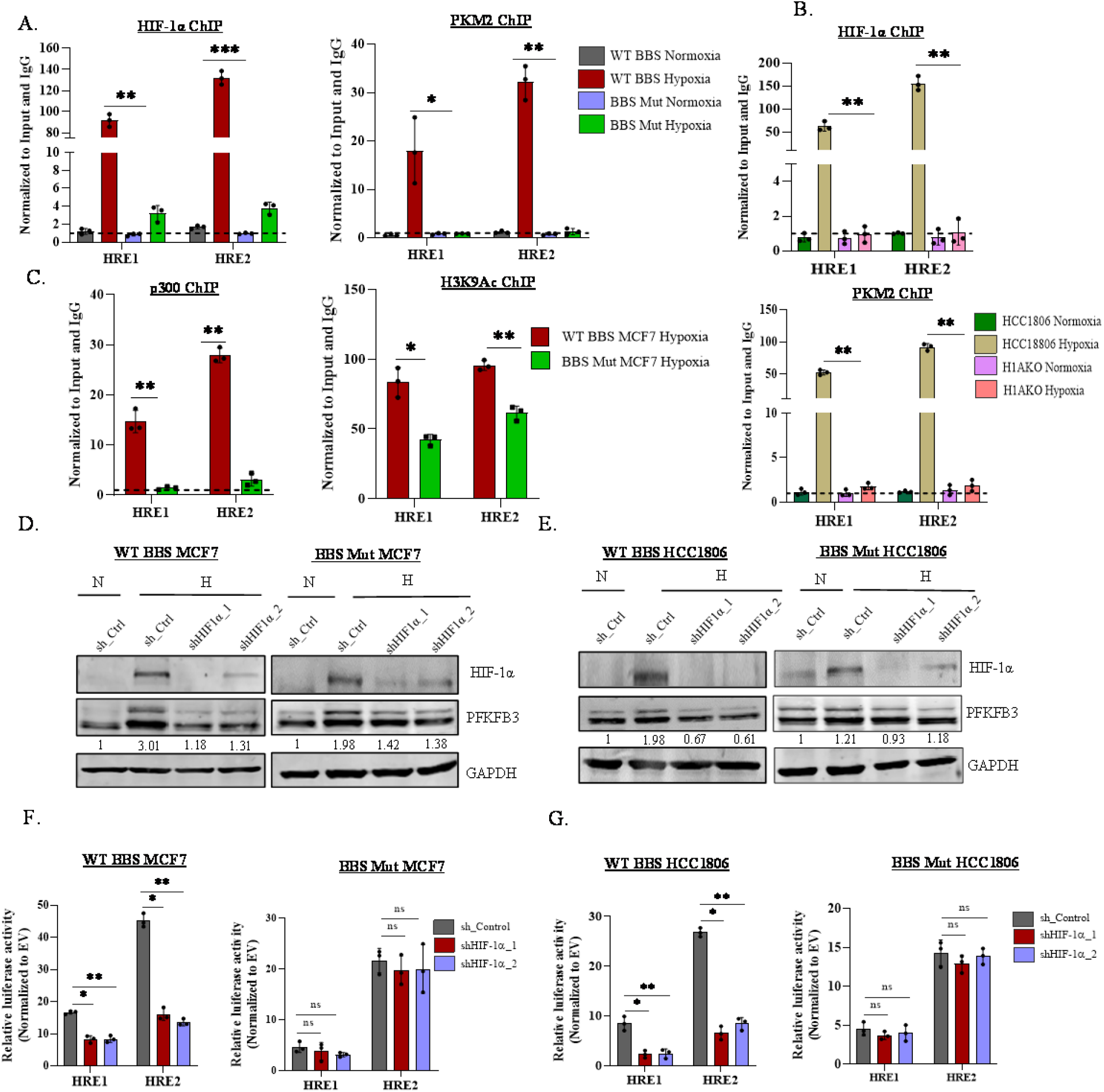
Hypoxic expression of PFKFB3 requires HIF-1α and PKM2. (A) ChIP assay of HIF-1α and PKM2 performed in WT BBS and BBS Mut MCF7 cells. (B) ChIP assay of HIF-1α and PKM2 in HCC1806 and HIAKO cells. Results show that PKM2 fails to occupy the *PFKFB3* HREs in the absence of HIF-1α. (C) ChIP assay of p300 and H3K9Ac in WT BBS and BBS Mut MCF7 cells under hypoxia. Immunoblot analysis of HIF-1α knockdown performed in WT BBS and BBS Mut cell lines of (D) MCF7 and (E) HCC1806. Luciferase assay performed in WT BBS and BBS Mut cell lines of (F) MCF7 and (G) HCC1806 shows the reduction in the FLuc activity of HRE1 and HRE2 only in the WT BBS cell lines upon performing HIF-1α knockdown. For (D), and (E) representative images are provided. Error bars show mean values ± SD (n = 3 unless otherwise specified). As calculated using two-tailed Student’s t-test, *P ≤ 0.05, **P ≤ 0.01, ***P ≤ 0.001, ****P ≤ 0.0001.

Interestingly, HIF-1α enrichment was not observed on HRE1 and HRE2 in BBS Mut MCF7 cells under hypoxia (Fig. 3A). To investigate if in the absence of PKM2, PFKFB3 regulation exhibits diminished dependency on HIF-1α, we performed HIF-1α knockdown. Results showed insignificant decrease of PFKFB3 expression only in the BBS Mut cell lines (Fig. 3D, Fig. 3E). Additionally, the FLuc activity corresponding to HRE1 and HRE2 was dampened only in the WT BBS cell lines upon performing HIF-1α knockdown (Fig. 3F, Fig. 3G). Conclusively, our observations suggest that PKM2 enhances binding of HIF-1α and p300 at *PFKFB3* HREs. PKM2 also is an essential interacting partner of HIF-1α, without which the transcription factor has reduced affinity towards *PFKFB3* HREs (Fig. 2E, 3A). Furthermore, *PFKFB3* is regulated by yet unreported transcription factors other than HIF-1α in the absence of PKM2 which maintains its basal-level expression.

### 5. HIF-2α acts as the master regulator of *PFKFB3* in the absence of HIF-1α or PKM2

HIF-1α and HIF-2α are known to recognize an identical DNA motif (41–43). However, HIF1 and HIF2 complex regulate different genes, and only a few shared targets have been identified (41, 44). To investigate the intriguing possibility if the reduced occupancy of HIF-1α rendered *PFKFB3* HREs available for HIF-2α binding, ChIP assay with HIF-2α antibody was performed. Interestingly, results indicated a prominent enrichment of HIF-2α on HRE1 and HRE2 in the BBS Mut MCF7 cells (Fig. 4A). Moreover, HIF-2α expression did not exhibit variation across all the WT and CRISPR/Cas9 mutants used throughout our study (Fig. S4.A). These findings suggest that HIF-2α is stabilized under hypoxia and its differential occupancy on *PFKFB3* HREs occurs due to reduced occupancy of HIF-1α triggered by absence of PKM2.

**Figure 4.**
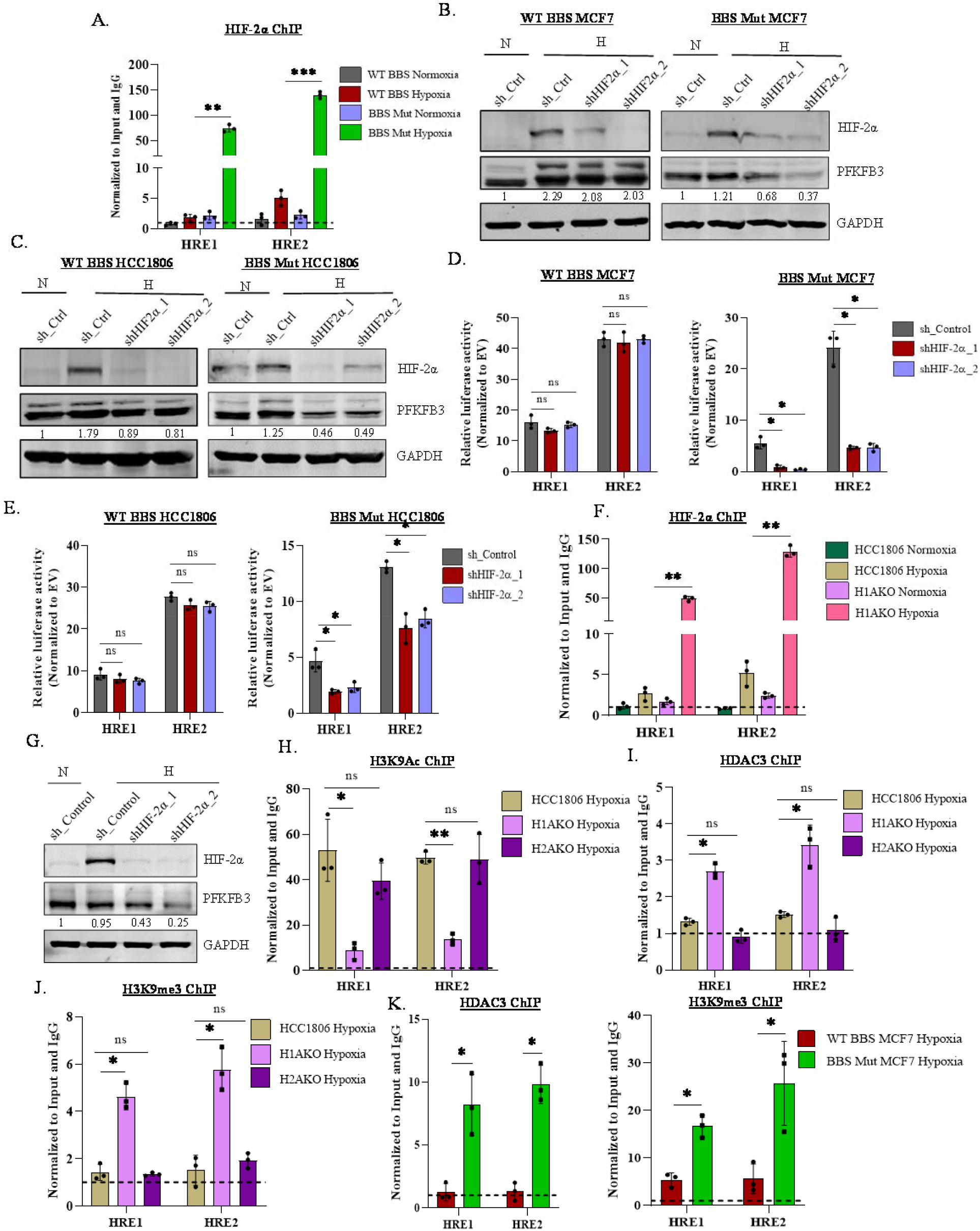
PFKFB3 is a novel non-canonical target of HIF-2α. (A) ChIP qRT-PCR analysis indicating a predominant occupancy of HIF-2α on PFKFB3 HREs under hypoxia in BBS Mut MCF7 cells. Immunoblot analysis of HIF-2α knockdown performed in WT BBS and BBS Mut cell lines of (B) MCF7 and (C) HCC1806. Luciferase activity of PFKFB3 HREs upon performing HIF-2α knockdown in WT BBS and BBS Mut cell lines of (D) MCF7 and (E) HCC1806 under hypoxia. (F) ChIP assay showing enrichment of HIF-2α on PFKFB3 HRE1 and HRE2 in H1AKO cells. (G) Immunoblot analysis of PFKFB3 upon performing HIF-2α knockdown in H1AKO cells. ChIP assay of (H) H3K9Ac, (I) HDAC3, and (J) H3K9me3 in HCC1806, H1AKO, and H2AKO cells under hypoxia. (K) ChIP assay of HDAC3 and H3K9Ac and H3K9me3 in WT BBS and BBS Mut MCF7 cells under hypoxia. For (B), (C), and (G), representative images are provided. Error bars show mean values ± SD (n = 3 unless otherwise specified). As calculated using two-tailed Student’s t-test, *P ≤ 0.05, **P ≤ 0.01, ***P ≤ 0.001, ****P ≤ 0.0001.

To further investigate this intriguing experimental observation, HIF-2α knockdown studies were performed. PFKFB3 was extensively downregulated only in the BBS Mut cell lines (Fig. 4B, Fig. 4C). Functional validation of the role of HIF-2α was also investigated using luciferase assay performed in the HIF-2α knockdown cells. We observed a significant decrease in the FLuc activity corresponding to HRE1 and HRE2 in BBS Mut cell lines. In contrast, there was an insignificant difference observed in the FLuc activity of the HREs in WT BBS cell lines (Fig. 4D, Fig. 4E).

Subsequently, to verify that PKM2 does not act as a coactivator of HIF-2α for maintaining PFKFB3 expression, we performed experiments using H1AKO cells. HIF-2α enrichment was also observed on HRE1 and HRE2 in H1AKO cells as indicated by ChIP assay results, highlighting a previously unreported opportunistic regulatory mechanism exhibited by HIF-2α (Fig. 4F). Furthermore, immunoblotting results evidently demonstrated that PFKFB3 expression was affected upon downregulating HIF-2α in H1AKO cells (Fig. 4G). As HIF-2α also interacts with HRE1 and HRE2, FLuc activity of these two sites was further dampened along with significant reduction of PFK activity upon eliminating HIF-2α in H1AKO cells (Fig. S4.B, S4.C). Additionally, we also generated CRISPR/Cas9-mediated HIF-2α knockout (H2AKO) cells of HCC1806 (Fig. S4.D). Upon performing PKM2 knockdown in H2AKO cells under hypoxia, we observed a hampered induction of PFKFB3 (Fig. S4.E).

Remarkably, our study shows that, unlike HIF-1α, HIF-2α does not cause extensive hypoxic induction of PFKFB3 and maintains its basal expression. To discern the underlying mechanism, we performed H3K9Ac ChIP assay using hypoxic HCC1806, H1AKO and H2AKO cells. The results indicated a loss of H3K9Ac marks at *PFKFB3* HREs only in H1AKO cells (Fig. 4H). Interestingly, despite of HIF-2α occupancy on *PFKFB3* HREs, there was decreased enrichment of p300 in H1AKO cells which suggested the involvement of another epigenetic factor preventing HIF-2α from optimally inducing *PFKFB3* (Fig. S4.G). To further explore the mechanism behind this phenomenon, we performed histone deacetylase 3 (HDAC3) ChIP assay in HCC1806, H1AKO and H2AKO cells. Our results clearly demonstrated enhanced presence of HDAC3 in H1AKO cells under hypoxia (Fig. 4I). Moreover, H1AKO cells also exhibited enhanced H3K9me3 histone modification (associated with gene repression) on *PFKFB3* HREs (Fig. 4J). We also performed HDAC3 and H3K9me3 ChIP assay in WT and BBS Mut MCF7 cells under hypoxia (Fig. 4K). The results clearly validated the vital contribution of the PKM2-HIF-1α complex in promoting p300-mediated epigenetic upregulation of *PFKFB3* under hypoxia. Consequently, absence of PKM2 results in HDAC3 recruitment at *PFKFB3* HREs which is responsible for the decreased H3K9Ac marks.

Additionally, we also observed an enrichment of histone deacetylase 3 (HDAC3) in H1AKO cells (Fig. S4.E) indicating that despite of HIF-2α occupancy, it is unable to efficiently induce PFKFB3 due to the diminished occupancy of p300 paired with enhanced HDAC3 activity. We also performed HDAC3 and H3K9me3 ChIP assay in WT and BBS Mut MCF7 cells under hypoxia. The results clearly validated that absence of PKM2 results in HDAC3 recruitment at *PFKFB3* HREs which is responsible for the decreased H3K9Ac marks (Fig. S4. F, Fig. S4. G).

Taken together, these data indicate that *PFKFB3* is a non-canonical target of HIF-2α as this transcription factor-mediated regulation prevails only in the absence of PKM2 or HIF-1α. Furthermore, as the differential occupancy of p300 and HDAC3 at *PFKFB3* HREs is orchestrated by PKM2, HIF-2α is unable to cause hypoxic induction of *PFKFB3*.

### 6. Shikonin inhibits the nuclear translocation of PKM2 to hamper hypoxic induction of PFKFB3

PKM2 is associated with various non-canonical functions that drive cancer progression and aggression; therefore, drugs targeting PKM2 present as desirable candidates for anti-cancer therapy. To investigate the effects of PKM2 inhibition under hypoxia, we employed the use of shikonin-a compound well-characterized to inhibit only PKM2 and not PKM1 (45). Molecular docking studies revealed that shikonin interacts with PKM2 with a best binding score of -8.1 kcal/mol. Additionally, the energy-optimized structure demonstrated that the inhibitor exclusively interacts with protein region coded by exon 10 (present only in PKM2) (Fig. S5.A). R399/400 residue has been previously reported to be involved in forming a putative nuclear localization signal (NLS) of PKM2 (46). Interestingly, the optimized poses of the protein and ligand interaction as analyzed by PyMOL 2.5 and subsequently visualized by PLIP indicated that R399/400 can form hydrogen bonds with shikonin (Fig. S5.B). Therefore, we suspected that subjecting hypoxic breast cancer cells to shikonin-mediated PKM2 inhibition will block its NLS and suppress non-canonical nuclear functions.

To test this hypothesis, we treated WT BBS cell lines with IC_50_ concentration of shikonin and exposed the cells to 24hrs of hypoxia (Fig. S5.C, S5.D). Immunofluorescence imaging performed to visualize the sub-cellular localization of PKM2 revealed its significantly reduced nuclear pool in the drug-treated cells (Fig. 5A, Fig. 5B). Furthermore, upon subjecting the cells to shikonin, only the WT BBS cell lines exhibited reduced hypoxic induction of PFKFB3, to indicate that shikonin inhibits the nuclear translocation of PKM2 (Fig. 5C, Fig. 5D) without altering its expression (Fig. S5.E). Additionally, we performed luciferase assay to verify the altered hypoxic induction of PFKFB3 in shikonin-treated WT BBS cell lines. The results indicated a decrease in the FLuc activity of HRE1 and HRE2 in both the WT BBS cell lines upon drug treatment. Since PFKFB3 is maintained in a PKM2-independent manner by HIF-2α, no significant difference was observed in the FLuc activity of the *PFKFB3* HREs in the BBS Mut cell lines (Fig. 5E, Fig. 5F). Consequently, the restricted PFKFB3 induction in the shikonin-treated WT BBS cell lines resulted in a decreased PFK activity, indicating that the rate of the committed step of glycolysis catalyzed by PFK-1 is affected (Fig. 5G).

**Figure 5.**
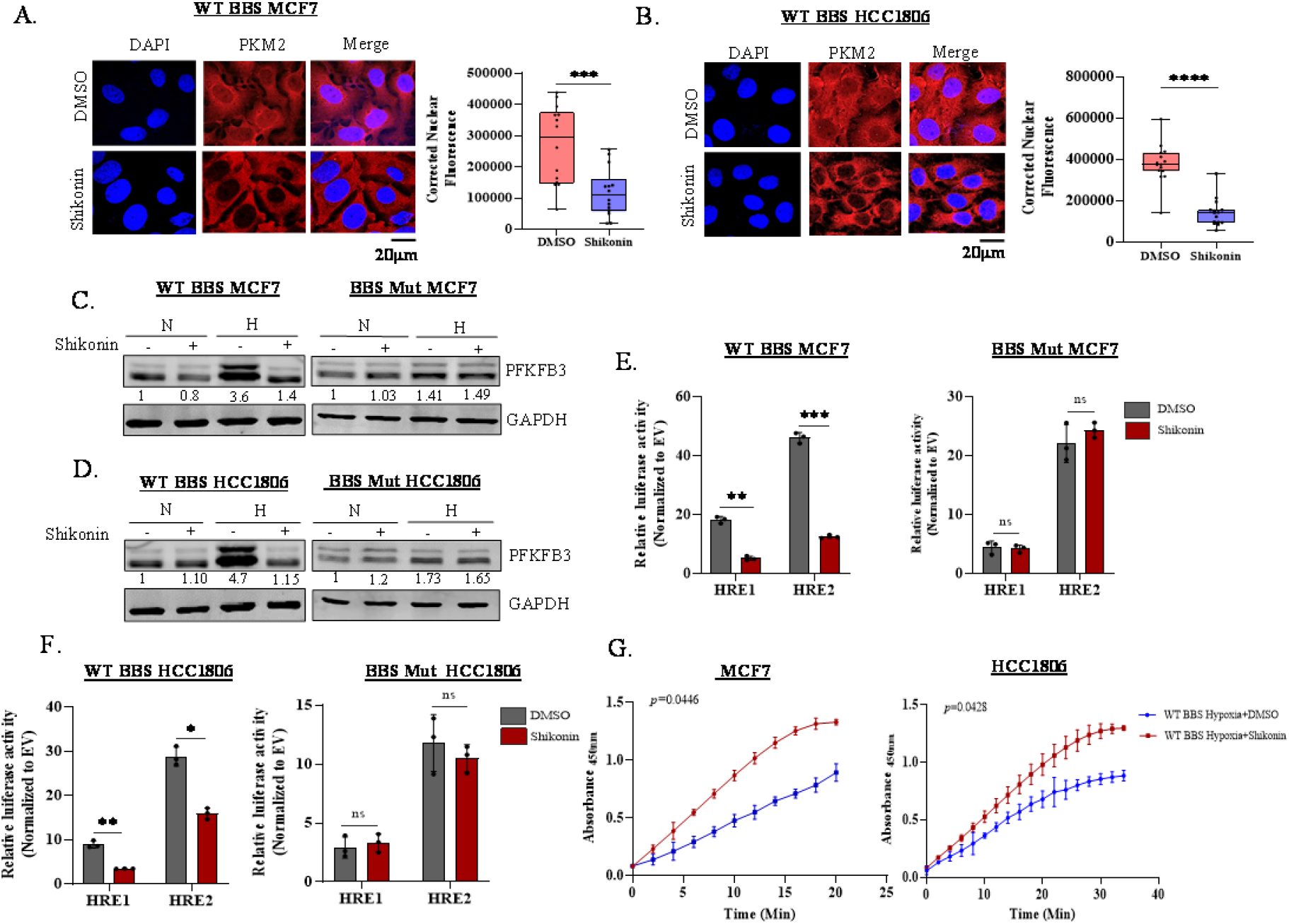
Shikonin treatment affects PFKFB3 expression in hypoxic breast cancer cells. Immunofluorescence imaging analysis and quantification showing blocked PKM2 nuclear translocation upon shikonin treatment in WT BBS (A) MCF7 and (B) HCC1806 cells. Magnification used: 60X. Immunoblot analysis of PFKFB3 expression upon treating WT BBS and BBS Mut cell lines of (C) MCF7 and (D) HCC1806 with shikonin for 24hrs under hypoxia. Luciferase assay performed upon subjecting shikonin treatment to the WT BBS and BBS Mut cells of (E) MCF7 and (F) HCC1806 cells under hypoxia. Results indicate a decreased activity of HRE1 and HRE2 in the WT BBS cell lines. (G) PFK Assay performed in WT BBS MCF7 and HCC1806 cells showing the reduced PFK activity upon shikonin treatment in hypoxia. For (A), (B), (C), (D) and (G), representative images are provided. Error bars show mean values ± SD (n = 3 unless otherwise specified). As calculated using two-tailed Student’s t-test, and unpaired t-test for PFK assay, *P ≤ 0.05, **P ≤ 0.01, ***P ≤ 0.001, ****P ≤ 0.0001.

Conclusively, our results evidently prove that PKM2-NLS interacting drugs can effectively restrict its nuclear translocation under hypoxia to inhibit PFKFB3 expression.

### 7. The PKM2-HIFs-PFKFB3 axis determines the glycolytic fate of hypoxic breast cancer cells

To explore our study’s clinical relevance and identify the effects of the PKM2-HIF-1α-mediated PFKFB3 regulation, we performed immunohistochemical analyses of tissue sections obtained from breast cancer patients. For identifying the hypoxic regions, tumor sections were probed with CAIX antibody along with PKM2 and PFKFB3. We found a strong co-expression of both the glycolytic enzymes in the CAIX-positive regions (Fig. 6A, Fig.6C). Additionally, the non-hypoxic regions (marked by negative staining for CAIX) exhibited reduced PKM2 and PFKFB3 expression, corroborating our findings that in hypoxic regions of breast cancers, PFKFB3 expression is primarily driven in a PKM2-HIF-1α-dependent manner (Fig. 6B, Fig. 6C).

**Figure 6.**
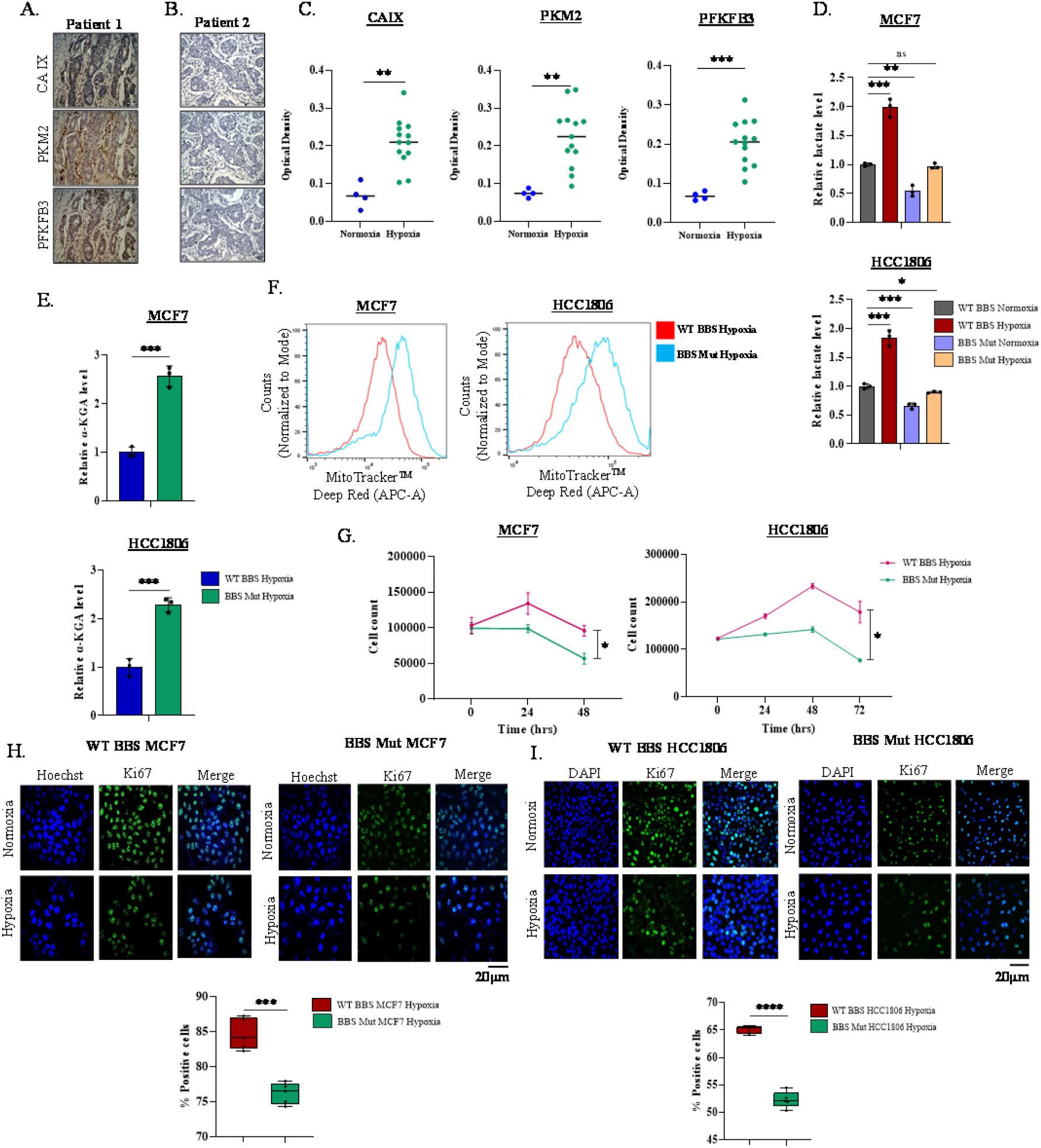
Functional significance of PKM2-HIFs-PFKFB3 axis. (A) Immunohistochemistry analysis of CAIX, PKM2, and PFKFB3. Hypoxic regions marked by the strong membranous and/or cytoplasmic immunostaining for CAIX also exhibit strong expression of PKM2 and PFKFB3. (B) Normoxic regions of tumors indicated by a negative staining for CAIX also shows relatively poor expression of PKM2 and PFKFB3. Magnification used: 40X. (C) Quantification of CAIX, PKM2 and PFKFB3 represented optical density for the normoxic and hypoxic regions of 18 breast cancer patients (D) Lactate production in the WT BBS and BBS Mut cell lines of MCF7 and HCC1806 cells assessed by lactate assay. (E) α-KGA assay indicating elevated α-KGA levels in the BBS Mut cell lines of MCF7 and HCC1806 under hypoxia. (F) MitoTracker™ Red dye FACS analysis post 24hrs of hypoxic treatment. Results show presence of relatively higher ΔΨ in BBS Mut cell lines. (G) Proliferation assay performed in WT BBS and BBS Mut cell lines of MCF7 and HCC1806 by subjecting the cells to the mentioned time period of 1% O_2_ treatment. Ki67 immunofluorescence imaging followed by quantification of % positive cells indicating a decreased proliferative potential of BBS Mut cell lines of (H) MCF7 and (I) HCC1806. Magnification used: 10X. For (A), (B), (F), (H), and (I), representative images are provided. Error bars show mean values ± SD (n = 3 unless otherwise specified). As calculated using two-tailed Student’s t-test, *P ≤ 0.05, **P ≤ 0.01, ***P ≤ 0.001, ****P ≤ 0.0001.

Lactate assay results indicated reduced intracellular lactate level in the BBS Mut cell lines as both PKM2 and PFKFB3 enhance the rate of glycolysis under hypoxia (Fig. 6D). Next, we determined if the decreased glycolysis caused by the loss of PKM2 and decreased PFKFB3 led to an increased dependence on oxidative phosphorylation (OXPHOS) for ATP production. To investigate this phenomenon, we estimated α-ketoglutarate (α-KGA) levels. We observed that the TCA cycle was more active in the PKM2 knockout cells as indicated by elevated α-KGA levels (Fig. 6E).

Additionally, direct evidence of an enhanced OXPHOS rate was also obtained by utilizing MitoTracker™ Deep Red, which is up-taken in mitochondrial membrane potential (ΔΨ) dependent manner (47, 48). WT BBS and BBS Mut cell lines cultured under 24hrs of hypoxia were treated with MitoTracker™ Deep Red dye and subjected to FACS analysis. The results demonstrated an enhanced accumulation of MitoTracker^™^ Deep Red in the BBS Mut cell lines (Fig. 6F). As the ΔΨ is positively correlated with enhanced mitochondrial activity (49), we performed mitochondrial complex I activity assay using mitochondria isolated from WT BBS and BBS Mut MCF7 cells. The purity of the mitochondrial isolate was verified by checking for cytoplasmic and nuclear protein markers (Fig. S6.A). In the assay, the absorbance at 600nm (as a function of time) is inversely proportional to the mitochondrial complex I activity. Our results indicated a rapid drop in the absorbance by the mitochondria isolated from hypoxic BBS Mut MCF7 to confirm the enhanced dependency on OXPHOS (Fig. S6.B).

### 8. Disrupting the PKM2-HIFs-PFKFB3 axis using shikonin hampers the proliferative potential of breast cancer cells

The metabolic state of a cancer cell is known to influence its replicative potential. A shift from a glycolysis-dependent state to an OXPHOS-dependent state is known to reduce the proliferation of cancer cells (50). Therefore, we monitored the proliferation rate of the WT BBS and BBS Mut cells upon subjecting them to hypoxic treatment. The BBS Mut cells replicated significantly slower than the WT BBS cells (Fig. 6G). Additionally, we also performed an Ki67 immunostaining and image analysis revealed higher % positive cells in the WT BBS cell lines than their respective BBS Mut counterpart cell lines (Fig. 6H, Fig. 6I).

Furthermore, to investigate the effects of blocking the PKM2-HIFs-PFKFB3 axis in an *in-vitro* model mimicking physiological conditions, we performed experiments using 3D-cultured breast cancer spheroids. The formation of spheroids often leads to the generation of hypoxic regions. We evaluated the presence of low O_2_ niches in the spheroids by utilizing Hypoxyprobe™-1 which forms stable covalent adducts with thiol groups in proteins, peptides, and amino acids under pO_2_<10mm Hg. As shown in Fig. S6.C, multiple hypoxic foci were detected in the dense spheroids formed by WT BBS HCC1806 cells. However, we could not detect hypoxic regions in WT BBS MCF7-derived spheroids probably because MCF7 cells are known to form loose spheroids (51, 52). Therefore, to understand the effects of inhibiting PKM2 under hypoxia, the subsequent 3D experiments were performed using WT BBS and BBS Mut HCC1806 cells.

All the cell lines were subjected to identical cultural conditions and were monitored for the changes in the spheroid size using bright field microscopy. The spheroid size plateaued post 7 days of seeding with an average diameter of 62±8.9μm for WT BBS HCC1806; whereas the BBS Mut HCC1806 formed remarkably smaller spheroids with a diameter size of 38.96±5.86μm to indicate a loss of proliferative potential of PKM2 knockout cell lines (Fig. 7A). Additionally, to understand the vitality of inhibiting PKM2 signalling under hypoxia, we treated the spheroids generated from WT BBS HCC1806 cells (post 5^th^ day of spheroid culture) with 2.2μM of shikonin for 48hrs. Microscopic analysis clearly indicated a significant decrease in the spheroid size post 48hrs of drug treatment (Fig. 7B). As we have demonstrated that shikonin blocks the nuclear transport of PKM2 to hamper PFKFB3 induction, we estimated that treating the 3D spheroids with the inhibitor would increase OXPHOS activity. Notably, we performed MitoTracker™ Deep Red staining and observed that shikonin treated spheroids showed increased uptake of MitoTracker™ Deep Red dye (Fig. S6.D).

**Figure 7.**
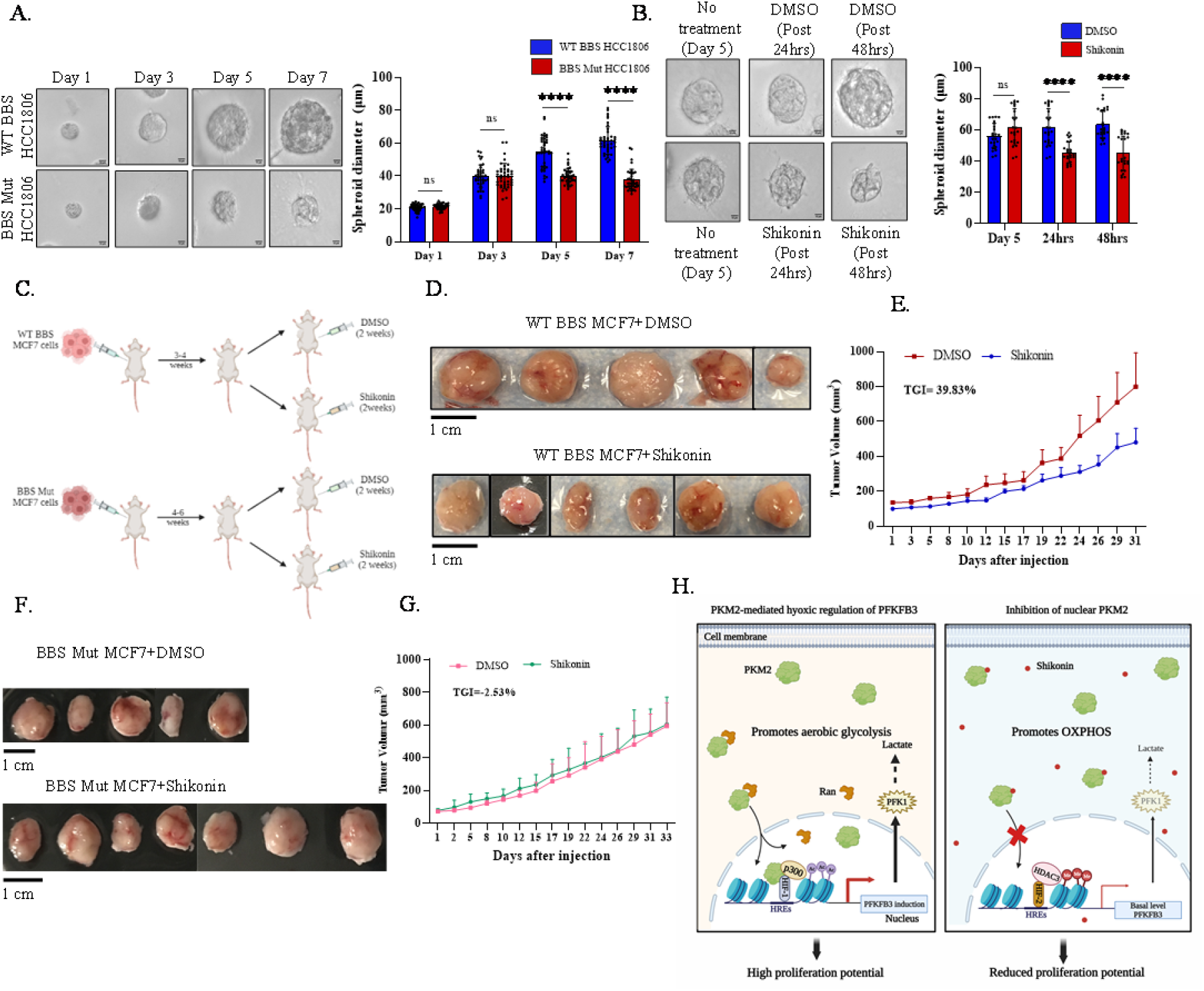
Shikonin blocks the HIFs-PKM2-PFKFB3 axis to exhibit anti-tumoral activity. (A) BBS Mut HCC1806 cells-derived 3D spheroids exhibit significantly less spheroid diameter compared to the WT BBS cells. Magnification used: 40X. (B) Shikonin treatment performed in WT BBS HCC1806-derived spheroids (post 5 days of growth). (C) Schematic representation of the *in-vivo* experimental protocol. (D) Representative tumor images from athymic nude mice bearing WT BBS MCF7 cells-derived tumors and receiving DMSO (n= 5) or shikonin (n= 6) treatment. (E) Line graph depicting relative growth rate of WT BBS MCF7 xenograft tumors in athymic nude mice treated either with DMSO or shikonin. Tumor growth inhibition (TGI) is indicated. (F) Representative tumor images from athymic nude mice bearing BBS Mut MCF7 cells-derived tumors and receiving DMSO (n= 5) or shikonin (n= 7) treatment. (G) Line graph depicting relative growth rate of BBS Mut MCF7 xenograft tumors in athymic nude mice treated either with DMSO or shikonin. Tumor growth inhibition (TGI) is indicated. (H) Model hypothesis. Hypoxia serves as a stimulus to promote the nuclear translocation of PKM2 in breast cancer cells. The nuclear PKM2 transcriptionally upregulates *PFKFB3* by enhancing binding of HIF-1α and p300. The enhanced PFKFB3 activity results in increased glycolytic flux and lactate production to promote aerobic glycolysis; contributing towards high proliferation rate. Restricting nuclear migration of PKM2 by eliminating Ran or treating the cancer cells with shikonin dampens the hypoxic induction of PFKFB3. In this case, a basal level of PFKFB3 expression is maintained by HIF-2α in a PKM2-independent manner. The lower PFKFB3 activity causes substantial inhibition of PFK1 to hamper glycolytic rate. Furthermore, this switch in the master transcriptional regulator of *PFKFB3* results in enhanced dependency on mitochondrial OXPHOS, which impedes the proliferative potential of hypoxic breast cancer cells. Notably, eliminating PKM2 enhances HDAC3 occupancy making PKM2 vital epigenetic regulator of PFKFB3 under hypoxia.

To verify if shikonin affects the proliferation of breast cancer cells *in-vivo*, we performed experiments using xenograft tumor models in athymic nude mice. Mice carrying WT BBS or BBS Mut MCF7 tumors were treated either with shikonin or DMSO and relative tumor growth was monitored over time (Fig. 7C). As evident from the tumor volume measured every alternate day for 4 weeks, shikonin treatment resulted in a marked decrease in WT BBS MCF7 tumor growth (Fig. 7D, Fig.7E). Moreover, there was no substantial difference observed between the tumor cell growth of BBS Mut cells receiving shikonin or DMSO treatment (Fig. 7F, Fig. 7G). Importantly, shikonin treatment exhibited >40-fold higher growth inhibitory effect on WT BBS MCF7 tumors (TGI= 39.83%) as compared to the BBS Mut tumors (TGI= -2.53%).

Taken in concert, the cell-based data as well as the *in-vivo* data consistently prove that inhibition of PKM2 severely hampers breast cancer cell proliferation by making them OXPHOS dependent.

## Discussion

It is estimated that almost 10 million cancer patient deaths occurred in 2020, and breast cancer was identified as the most commonly diagnosed type of cancer (53). Due to the multinodular structure of breast cancers, numerous hypoxic core regions surrounded by a normoxic frame of cells can often be detected (54). This microenvironmental condition of low pO_2_ is known to increase aggressiveness of cancers that ultimately presents as major hurdle in anti-cancer therapy (55, 56). An eminent feature of hypoxic breast cancer cells which presents as one of the paramount orchestrators of rewired cellular signalling is altered metabolism (1). Moreover, it is observed that the hypoxic niches also promote various non-canonical functions of metabolic enzymes to aid cancer cell survival (57). PKM2 is one such metabolic enzyme known to exhibit moonlighting functions under hypoxia and is also a critical contributor of the Warburg effect. The PKM2 isoform is preferentially expressed over PKM1 by cancer cells and undergoes nuclear translocation under hypoxia to transcriptionally regulate the expression of various genes. However, till date, the role of nuclear PKM2 in modulating global transcriptome and specially the expression of metabolism-related genes remains elusive primarily due to the difficulty in generating proper model systems which are selectively PKM2 knockouts (sparing the PKM1 isoform) in the context of cancer.

In order to specifically dissect the contributions of PKM2 in shaping hypoxic transcriptome, we compared the ChIP-seq analysis of HIF-1α with transcriptomic data obtained from comparing WT BBS and BBS Mut cells. Besides governing glycolytic rate, we identified for the first time, plethora of cellular pathways that are affected by PKM2; highlighting its role in sculpting global gene transcription. Additionally, Ran was identified to be the mediator of PKM2 translocation across the nuclear membrane under hypoxia. Transcriptome array analysis of the hypoxic WT BBS and BBS Mut MCF7 cells revealed *PFKFB3* as one of the top targets. Interestingly, we observed PKM2 localization on the *PFKFB3* HREs along with HIF-1α, indicating that the PKM2-HIF-1α interaction dominantly regulates hypoxic PFKFB3 induction. Among the three HREs, the HREs 1 and 2 were identified to be involved in mediating the transcriptional upregulation of *PFKFB3* in response to hypoxia, as evident from the ChIP and luciferase assays. Moreover, PKM2 directly recruited p300 and prevented HDAC3 from binding at *PFKFB3* HREs.

Another the key finding of our study is that in the absence of PKM2 or HIF-1α, PFKFB3 expression was not abrogated, but was comparable to its normoxic expression. This observation suggested the involvement of a potentially unreported transcription factor involved in governing the hypoxic regulation of PFKFB3 in the absence of HIF-1α or PKM2. Feeding on the fact that HIF-2α is also highly expressed under hypoxia and shares common binding sites with HIF-1α, we tested if HIF-2α had any contribution in maintaining basal expression of PFKFB3. Interestingly, we observed a significant enrichment of HIF-2α on the *PFKFB3* HREs in the HIF-1α and PKM2 knockout cells. *PFKFB3*, being non-canonical target of HIF-2α, gets weakly induced, ultimately aiding the cancer cells to maintain a basal level expression of the glycolytic enzyme in a PKM2-independent manner (Fig. 4A-Fig. 4C). As PFKFB3 is a critical regulator of glycolysis, our study discloses a unique epigenetic mechanism adapted by hypoxic cancer cells to ensure the continuity of glycolysis by maintaining PFKFB3 expression in the absence of its master regulators PKM2 and HIF-1α. Moreover, to the best of our knowledge, our investigations for the first time, provide detailed experimental insights into hypoxic epigenetic regulation of PFKFB3.

Notably, our study shows PKM2-independent transcriptional regulation of *PFKFB3* by HIF-2α results in a shift from glycolysis to mitochondrial OXPHOS under hypoxia. Mitochondrial respiration can function even at 0.5% of O_2_ availability (58, 59). Luengo and co-workers have previously negatively correlated ΔΨ with cellular proliferation (50). Consistently, we also observed a decrease in the proliferation of the BBS Mut cell lines, which are predominantly OXPHOS-dependent. Finally, we investigate the clinical relevance of this study by employing PKM2 inhibitor-shikonin. Shikonin dampens the hypoxic induction of PFKFB3 by interacting with a putative-NLS of PKM2 and preventing its hypoxic nuclear translocation. Notably, this phenomenon was coupled with a concomitant rise in the ΔΨ, a direct indicator of OXPHOS rate as verified using 2D and 3D cell culture methods. Moreover, mice carrying WT BBS MCF7 tumors and subjected to shikonin treatment exhibited remarkably decreased tumor growth, highlighting the paramount importance of targeting PKM2 signaling to impede tumor progression.

Evidently, our findings delineate a unique regulatory network that exists between two crucial glycolytic enzymes-PKM2 and PFKFB3 under hypoxia (Fig.7H). To the best of our knowledge, our work also highlights for the first time a novel phenomenon in which HIF-2α compensates for the reduced occupancy of HIF-1α to regulate *PFKFB3*. Furthermore, it will be interesting to identify the epigenetic writer negatively affected by PKM2 binding causes H3K9me3 enrichment at *PFKFB3* HREs. We further identify that disrupting this reported axis can lead to the impaired proliferation of hypoxic breast cancer cells. As PKM2 and PFKFB3 are critical enzymes enhancing glycolysis under hypoxia, implementing PKM2-targeting drugs as a part of anti-cancer regimen can be beneficial in combating the adverse repercussions of hypoxic cancer cells by exerting suppressive effects on the non-canonical functions of PKM2.

## Authors’ Contributions

S.S and M.R.P conceived and designed the study. M.R.P contributed to ChIP, immunoblotting, Co-IP, luciferase, qRT-PCR assays, 3D-spheroid experiments, FACS, knockdown and CRISPR/Cas9 experiments as well as generation of schematic representations for the manuscript. M.R.P and S.A.M contributed to performing HTA 2.0 array. ChIP-seq and HTA 2.0 array data was analyzed by A.R. M.R.P and A.R contributed in performing metabolic assays. M.R.P and S.A.M contributed to proliferation assays, confocal microscopy and image analyses. A.R contributed to molecular docking studies and cloning of the used plasmid constructs. S.A.M contributed to immunohistochemical analysis of patient-derived tumor sections. K.B contributed to animal experiments. M.R.P, A.R, and S.S analyzed the data. A.S, J.M, S.K.S and S.S contributed new reagents. S.S, S.K.S, and M.R.P contributed to scientific discussions. M.R.P wrote the manuscript with input from all authors. S.S oversaw all the experiments, data analyses, and manuscript preparation.

## Acknowledgments

Graphical representations are generated using BioRender. This work is supported by Wellcome Trust/Department of Biotechnology (DBT) India Alliance Fellowship Grant IA/I/16/2/502719 (to S. Shukla). University Grants Commission (to M.R.P); Indian Institute of Science Education and Research Bhopal fellowship (to S.A.M) and Department of Science and Technology, Ministry of Science and Technology (to A.R).

## References

1. D. Hanahan, R. A. Weinberg, Hallmarks of cancer: The next generation. Cell (2011) https://doi.org/10.1016/j.cell.2011.02.013.

2. P. P. Hsu, D. M. Sabatini, Cancer cell metabolism: Warburg and beyond. Cell 134, 703–707 (2008).

3. C. Chen, N. Pore, A. Behrooz, F. Ismail-Beigi, A. Maity, Regulation of glut1 mRNA by hypoxia-inducible factor-1: Interaction between H-ras and hypoxia. J. Biol. Chem. 276, 9519–9525 (2001).

4. G. L. Semenza, P. H. Roth, H. M. Fang, G. L. Wang, Transcriptional regulation of genes encoding glycolytic enzymes by hypoxia-inducible factor 1. J. Biol. Chem. (1994).

5. W. G. Kaelin, P. J. Ratcliffe, Oxygen Sensing by Metazoans: The Central Role of the HIF Hydroxylase Pathway. Mol. Cell 30, 393–402 (2008).

6. M. Ohh, et al., Ubiquitination of hypoxia-inducible factor requires direct binding to the β-domain of the von Hippel - Lindau protein. Nat. Cell Biol. 2, 423–427 (2000).

7. E. Ikeda, M. G. Achen, G. Breier, W. Risau, Hypoxia-induced transcriptional activation and increased mRNA stability of vascular endothelial growth factor in C6 glioma cells. J. Biol. Chem. 270, 19761–19766 (1995).

8. G. L. Semenza, et al., Hypoxia response elements in the aldolase A, enolase 1, and lactate dehydrogenase a gene promoters contain essential binding sites for hypoxia-inducible factor 1. J. Biol. Chem. (1996) https://doi.org/10.1074/jbc.271.51.32529.

9. R. H. Wenger, A. Rolfs, H. H. Marti, C. Bauer, M. Gassmann, Hypoxia, a novel inducer of acute phase gene expression in a human hepatoma cell line. J. Biol. Chem. 270, 27865–27870 (1995).

10. S. Amin, P. Yang, Z. Li, Pyruvate kinase M2: A multifarious enzyme in non-canonical localization to promote cancer progression. Biochim. Biophys. Acta - Rev. Cancer (2019) https://doi.org/10.1016/j.bbcan.2019.02.003.

11. N. Azoitei, et al., PKM2 promotes tumor angiogenesis by regulating HIF-1aα through NF-κB activation. Mol. Cancer 15, 1–15 (2016).

12. W. Luo, et al., Pyruvate kinase M2 is a PHD3-stimulated coactivator for hypoxia-inducible factor 1. Cell 145, 732–744 (2011).

13. S. Singh, et al., Intragenic DNA methylation and BORIS-mediated cancer-specific splicing contribute to the Warburg effect. Proc. Natl. Acad. Sci. U. S. A. (2017) https://doi.org/10.1073/pnas.1708447114.

14. N. Ahuja, et al., Hypoxia-induced TGF-β–RBFOX2–ESRP1 axis regulates human MENA alternative splicing and promotes EMT in breast cancer. NAR Cancer 2, 1–17 (2020).

15. J. Schödel, et al., High-resolution genome-wide mapping of HIF-binding sites by ChIP-seq. Blood 117, e207–e217 (2011).

16. S. X. Ge, D. Jung, D. Jung, R. Yao, ShinyGO: a graphical gene-set enrichment tool for animals and plants. Bioinformatics 36, 2628–2629 (2020).

17. Z. Zhou, et al., Oncogenic kinase–induced PKM2 tyrosine 105 phosphorylation converts nononcogenic PKM2 to a tumor promoter and induces cancer stem–like cells. Cancer Res. 78, 2248–2261 (2018).

18. C. Frezza, S. Cipolat, L. Scorrano, Organelle isolation: Functional mitochondria from mouse liver, muscle and cultured filroblasts. Nat. Protoc. 2, 287–295 (2007).

19. O. Trott, A. J. Olson, AutoDock Vina: Improving the speed and accuracy of docking with a new scoring function, efficient optimization, and multithreading. J. Comput. Chem., NA-NA (2009).

20. J. D. Dombrauckas, B. D. Santarsiero, A. D. Mesecar, Structural basis for tumor pyruvate kinase M2 allosteric regulation and catalysis. Biochemistry 44, 9417–9429 (2005).

21. H. M. Berman, et al., The Protein Data Bank. Nucleic Acids Res. 28, 235–242 (2000).

22. E. H. Baugh, S. Lyskov, B. D. Weitzner, J. J. Gray, Real-time PyMOL visualization for Rosetta and PyRosetta. PLoS One 6 (2011).

23. E. Krieger, et al., Improving physical realism, stereochemistry and side-chain accuracy in homology modeling: four approaches that performed well in CASP8. Proteins 77, 114 (2009).

24. M. F. Adasme, et al., PLIP 2021: Expanding the scope of the protein-ligand interaction profiler to DNA and RNA. Nucleic Acids Res. 49, W530–W534 (2021).

25. J. Debnath, S. K. Muthuswamy, J. S. Brugge, Morphogenesis and oncogenesis of MCF-10A mammary epithelial acini grown in three-dimensional basement membrane cultures. Methods 30, 256–268 (2003).

26. M. Vaapil, et al., Hypoxic Conditions Induce a Cancer-Like Phenotype in Human Breast Epithelial Cells. PLoS One 7 (2012).

27. B. Ducommun, et al., Abstract 4404: Multicellular tumor spheroid models to evaluate drugs targeting cell cycle checkpoints in 3D. 4404–4404 (2013).

28. M. C. Hsu, et al., Extracellular PKM2 induces cancer proliferation by activating the EGFR signaling pathway. Am. J. Cancer Res. 6, 628–638 (2016).

29. L. Li, Y. Zhang, J. Qiao, J. J. Yang, Z. R. Liu, Pyruvate kinase M2 in blood circulation facilitates tumor growth by promoting angiogenesis. J. Biol. Chem. 289, 25812–25821 (2014).

30. J. Liang, et al., Mitochondrial PKM2 regulates oxidative stress-induced apoptosis by stabilizing Bcl2. Cell Res. 27, 329–351 (2017).

31. L. Lv, et al., Mitogenic and Oncogenic Stimulation of K433 Acetylation Promotes PKM2 Protein Kinase Activity and Nuclear Localization. Mol. Cell (2013) https://doi.org/10.1016/j.molcel.2013.09.004.

32. Y. C. Yang, et al., Nuclear translocation of PKM2/AMPK complex sustains cancer stem cell populations under glucose restriction stress. Cancer Lett. 421, 28–40 (2018).

33. N. Hay, Reprogramming glucose metabolism in cancer: Can it be exploited for cancer therapy? Nat. Rev. Cancer (2016) https://doi.org/10.1038/nrc.2016.77.

34. L. Lu, Y. Chen, Y. Zhu, The molecular basis of targeting PFKFB3 as a therapeutic strategy against cancer. Oncotarget 8, 62793–62802 (2017).

35. L. Shi, H. Pan, Z. Liu, J. Xie, W. Han, Roles of PFKFB3 in cancer. Signal Transduct. Target. Ther. (2017) https://doi.org/10.1038/sigtrans.2017.44.

36. A. La Belle Flynn, et al., Autophagy inhibition elicits emergence from metastatic dormancy by inducing and stabilizing Pfkfb3 expression. Nat. Commun. 10, 1–15 (2019).

37. H. M. Li, et al., Blockage of glycolysis by targeting PFKFB3 suppresses tumor growth and metastasis in head and neck squamous cell carcinoma. J. Exp. Clin. Cancer Res. 36, 1–12 (2017).

38. F. Peng, et al., PFKFB3 is involved in breast cancer proliferation, migration, invasion and angiogenesis. Int. J. Oncol. 52, 945–954 (2018).

39. M. Obach, et al., 6-Phosphofructo-2-kinase (pfkfb3) gene promoter contains hypoxia-inducible factor-1 binding sites necessary for transactivation in response to hypoxia. J. Biol. Chem. (2004) https://doi.org/10.1074/jbc.M406096200.

40. S. A. Dames, M. Martinez-Yamout, R. N. De Guzman, H. Jane Dyson, P. E. Wright, Structural basis for Hif-1α/CBP recognition in the cellular hypoxic response. Proc. Natl. Acad. Sci. U. S. A. (2002) https://doi.org/10.1073/pnas.082121399.

41. V. L. Dengler, M. D. Galbraith, J. M. Espinosa, Transcriptional regulation by hypoxia inducible factors. Crit. Rev. Biochem. Mol. Biol. (2014) https://doi.org/10.3109/10409238.2013.838205.

42. J. W. Lee, J. Ko, C. Ju, H. K. Eltzschig, Hypoxia signaling in human diseases and therapeutic targets. Exp. Mol. Med. 51, 1–13 (2019).

43. H. Tian, S. L. McKnight, D. W. Russell, Endothelial PAS domain protein 1 (EPAS1), a transcription factor selectively expressed in endothelial cells. Genes Dev. 11, 72–82 (1997).

44. N. Skuli, et al., Endothelial HIF-2α regulates murine pathological angiogenesis and revascularization processes. J. Clin. Invest. 122, 1427–1443 (2012).

45. J. Chen, et al., Shikonin and its analogs inhibit cancer cell glycolysis by targeting tumor pyruvate kinase-M2. Oncogene 30, 4297–4306 (2011).

46. W. Yang, et al., ERK1/2-dependent phosphorylation and nuclear translocation of PKM2 promotes the Warburg effect. Nat. Cell Biol. 14, 1295–1304 (2012).

47. A. W. Greene, et al., Mitochondrial processing peptidase regulates PINK1 processing, import and Parkin recruitment. EMBO Rep. 13, 378–385 (2012).

48. R. Zhou, A. S. Yazdi, P. Menu, J. Tschopp, A role for mitochondria in NLRP3 inflammasome activation. Nature 469, 221–226 (2011).

49. I. Martínez-reyes, et al., for Diverse Biological Functions. Mol Cell 61, 199–209 (2016).

50. A. Luengo, et al., Increased demand for NAD+ relative to ATP drives aerobic glycolysis. Mol. Cell (2021) https://doi.org/10.1016/j.molcel.2020.12.012.

51. Y. Imamura, et al., Comparison of 2D- and 3D-culture models as drug-testing platforms in breast cancer. Oncol. Rep. 33, 1837–1843 (2015).

52. C. Te Kuo, et al., Three-dimensional spheroid culture targeting versatile tissue bioassays using a PDMS-based hanging drop array. Sci. Rep. 7, 1–10 (2017).

53. H. Sung, et al., Global Cancer Statistics 2020: GLOBOCAN Estimates of Incidence and Mortality Worldwide for 36 Cancers in 185 Countries. CA. Cancer J. Clin. 71, 209–249 (2021).

54. M. R. Pandkar, S. G. Dhamdhere, S. Shukla, Oxygen gradient and tumor heterogeneity: The chronicle of a toxic relationship. Biochim. Biophys. Acta - Rev. Cancer 1876, 188553 (2021).

55. P. Vaupel, A. Mayer, Hypoxia in cancer: Significance and impact on clinical outcome. Cancer Metastasis Rev. (2007) https://doi.org/10.1007/s10555-007-9055-1.

56. W. R. Wilson, M. P. Hay, Targeting hypoxia in cancer therapy. Nat. Rev. Cancer (2011) https://doi.org/10.1038/nrc3064.

57. D. Williams, B. Fingleton, Non-canonical roles for metabolic enzymes and intermediates in malignant progression and metastasis. Clin. Exp. Metastasis (2019) https://doi.org/10.1007/s10585-019-09967-0.

58. N. S. Chandel, G. R. S. Budinger, P. T. Schumacker, Molecular oxygen modulates cytochrome c oxidase function. J. Biol. Chem. 271, 18672–18677 (1996).

59. W. L. Rumsey, C. Schlosser, E. M. Nuutinen, M. Robiolio, D. F. Wilson, Cellular energetics and the oxygen dependence of respiration in cardiac myocytes isolated from adult rat. J. Biol. Chem. 265, 15392–15399 (1990).

