## Supplementary Figures and Tables for "PKM2-mediated epigenetic reprogramming regulates hypoxic expression of *PFKFB3* to promote breast cancer progression"

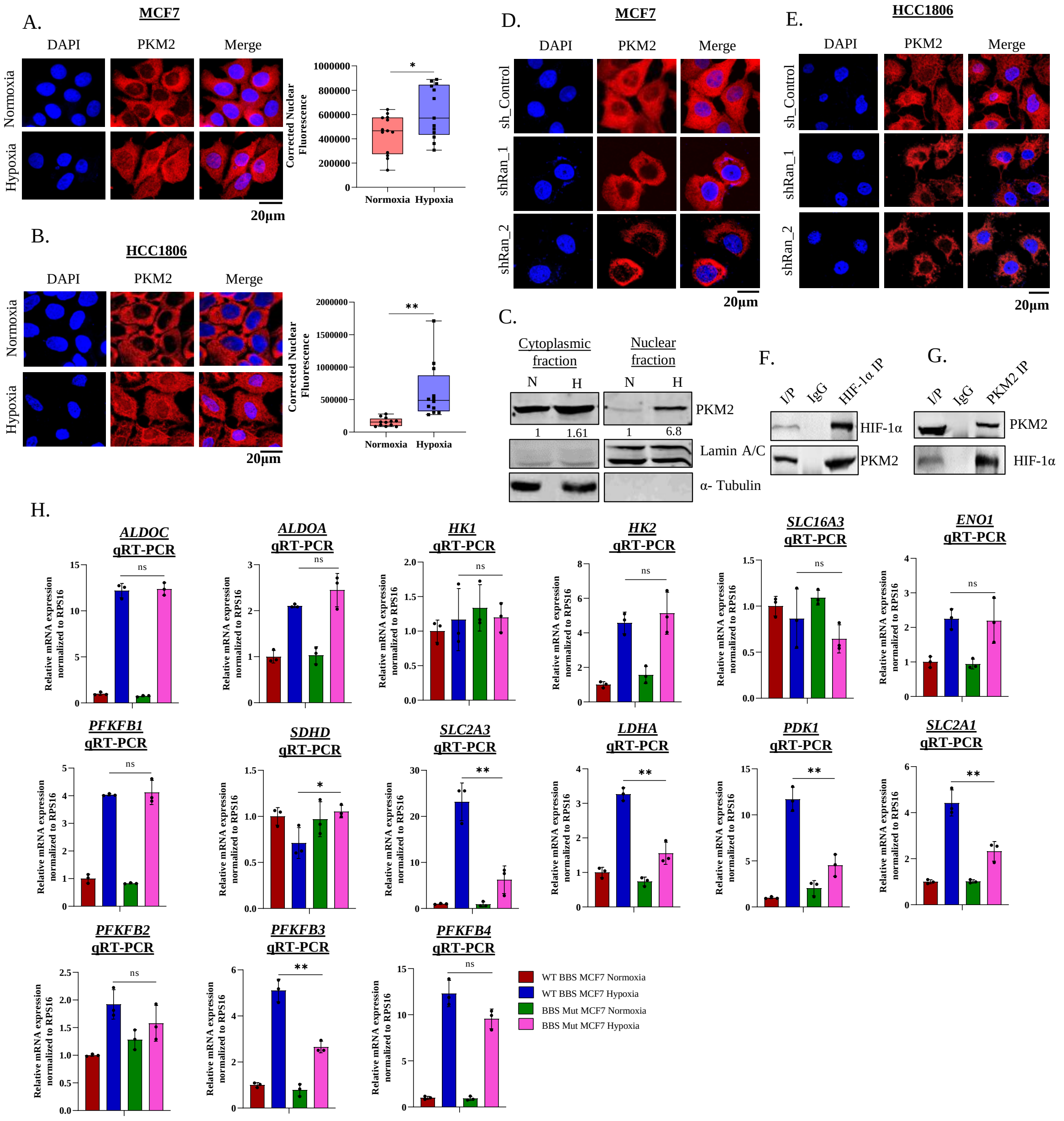


**Figure S1: PKM2 exhibits nuclear non-canonical function under hypoxia.** Immunofluorescence imaging and quantification performed under normoxia and hypoxia in (A) MCF7 and (B) HCC1806 indicating enriched pool of nuclear PKM2 under hypoxia. Magnification used: 60X. (C) Immunoblot analysis of the nuclear and cytoplasmic protein fractions isolated from normoxic and hypoxic HCC1806 cells. Immunofluorescence imaging showing hampered nuclear translocation of PKM2 upon performing Ran knockdown in (D) MCF7 and (E) HCC1806 cell lines under hypoxia. Magnification used: 60X. (F) and (G) Co-immunoprecipitation assay performed in MCF7 cells under hypoxia indicating the physical interaction of HIF-1α and PKM2. (H) qRT-PCR analysis of glycolytic gene targets of HIF-1α in WT BBS and BBS Mut MCF7 cells under hypoxia. For (A), (B), (C), (D), (E), (F) and (G), representative images are provided. Error bars show mean values ± SD (n = 3 unless otherwise specified). As calculated using two-tailed Student's t-test, $^{*}P \le 0.05$, $^{**}P \le 0.01$, $^{***}P \le 0.001$, $^{****}P \le 0.0001$.

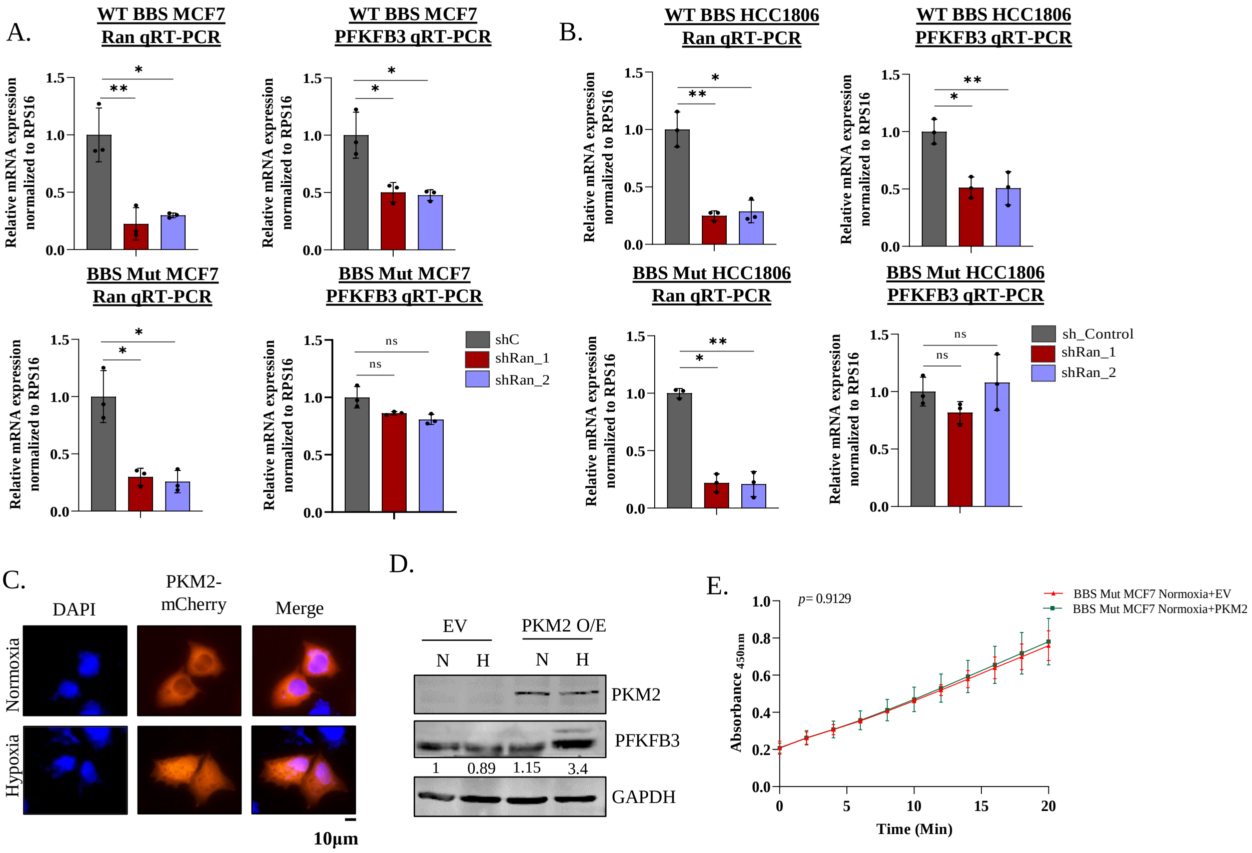


**Figure S2: Ran aids nuclear translocation of PKM2 to enhance PFKFB3 expression.** RPS16 normalized qRT-PCR analysis of *PFKFB3* in WT BBS and BBS Mut cell lines of (A) MCF7 (B) and HCC1806 cells upon performing Ran knockdown under hypoxia. Ran knockdown reduces hypoxic induction of *PFKFB3* in the WT BBS cell lines by blocking nuclear translocation of PKM2. (C) mCherry tagged PKM2 overexpression followed by fluorescence imaging showed enhanced nuclear translocation of PKM2 under hypoxia in the BBS Mut MCF7 cells. Magnification used: 40X. (D) Immunoblotting results depicting a rescued PFKFB3 expression in BBS Mut MCF7 cells upon PKM2 overexpression under hypoxia. (E) PFK assay showing no effect of PKM2 over-expression under normoxic condition on the overall PFK activity of BBS Mut MCF7 cells. Error bars show mean values ± SD (n = 3 unless otherwise specified). For (C), (D), (E), representative images are provided. As calculated using two-tailed Student's t-test, and unpaired t-test for PFK assay, $*P \leq 0.05$, $**P \leq 0.01$, $***P \leq 0.001$, $****P \leq 0.0001$.

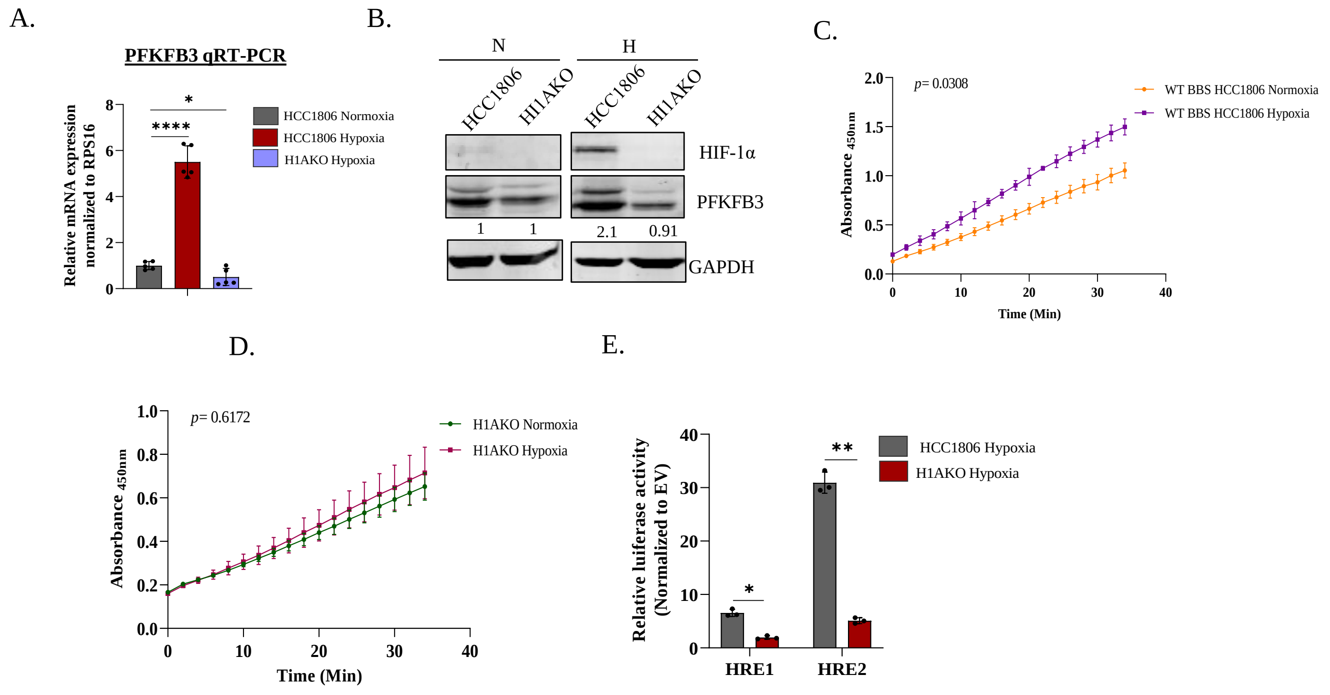


**Figure S3: HIF-1α is the master-regulator of PFKFB3 under hypoxia.** (A) RPS16 normalized qRT-PCR data showing reduced PFKFB3 expression in H1AKO cells. (B) Immunoblot analysis of PFKFB3 expression in HCC1806 and H1AKO. (C) PFK assay performed in WT BBS HCC1806 showing enhanced PFK activity under hypoxia. (D) PFK assay performed in H1AKO showing lack of significant enhancement of the PFK activity under hypoxia. (E) Luciferase assay results demonstrating reduced FLuc activity of *PFKFB3* HRE1 and HRE2 in H1AKO cells under hypoxia. For (B), (C) and (D), representative images are provided. Error bars show mean values ± SD (n = 3 unless otherwise specified). As calculated using two-tailed Student's t-test and unpaired t-test for PFK assay, $*P \leq 0.05$, $**P \leq 0.01$, $***P \leq 0.001$, $****P \leq 0.0001$.

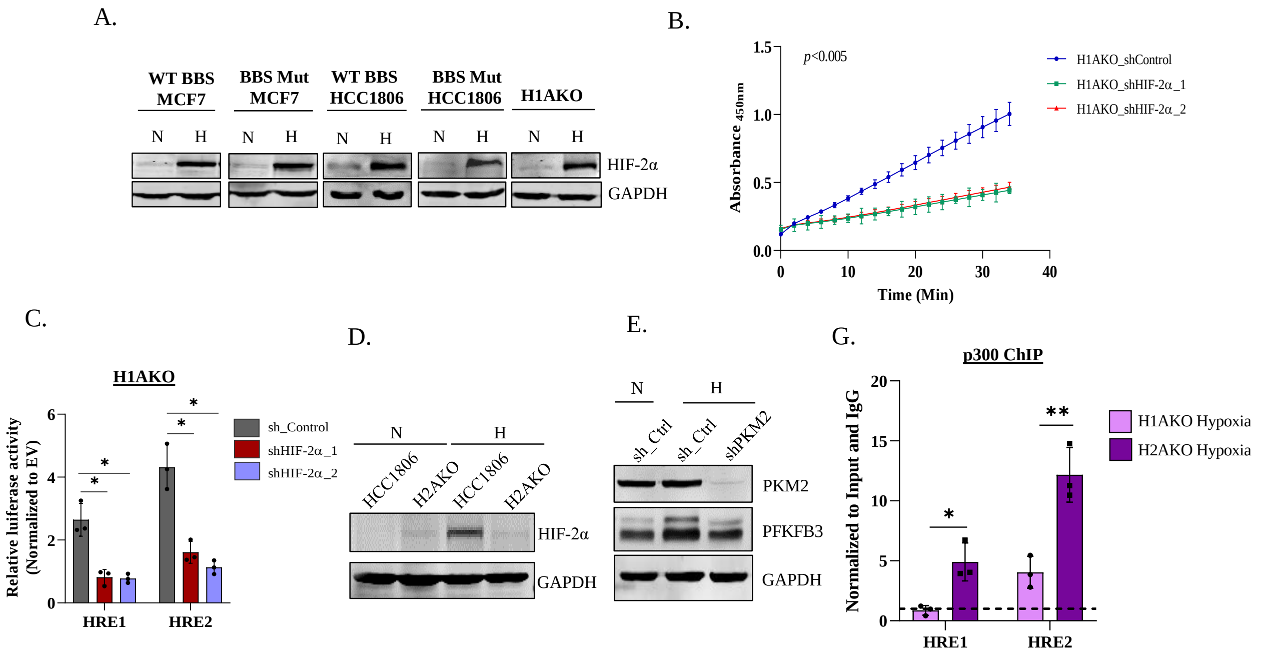


**Figure S4: HIF-2α maintains a basal-level expression of PFKFB3 in absence of HIF-1α or PKM2.** (A) Immunoblot analysis of HIF-2α in various WT and mutant cell lines used in study. (B) PFK Assay performed in the HIF-2α knockdown cells of H1AKO indicating a drastic reduction of PFK activity compared to the control cells under hypoxia. (C) Luciferase assay depicting the HIF-2α mediated transactivation of *PFKFB3* by showing the reduced functionality of the HREs upon performing HIF-2α knockdown in H1AKO cells. (D) Immunoblot analysis of H2AKO cells. (E) Western blot analysis showing decrease in the PFKFB3 upon performing PKM2 knockdown in H2AKO cells. (F) ChIP assay of p300 in H1AKO and H2AKO cells under hypoxia. For (A), (B), (D), and (E), representative images are provided. Error bars show mean values ± SD (n = 3 unless otherwise specified). As calculated using two-tailed Student’s t-test, and unpaired t-test for PFK assay $*P \leq 0.05$, $**P \leq 0.01$, $***P \leq 0.001$, $****P \leq 0.0001$.

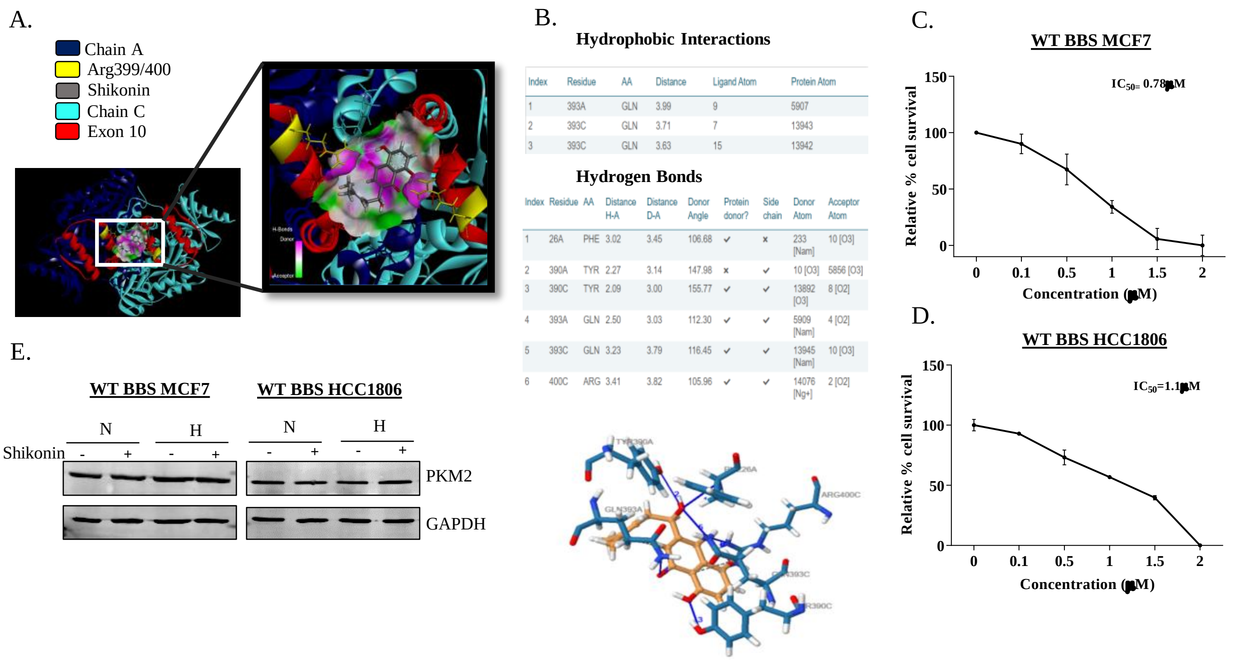


**Hydrophobic Interactions**

| Index | Residue | AA | Distance | Ligand Atom | Protein Atom |
|---|---|---|---|---|---|
| 1 | 393A | GLN | 3.99 | 9 | 5907 |
| 2 | 393C | GLN | 3.71 | 7 | 13943 |
| 3 | 393C | GLN | 3.63 | 15 | 13942 |

**Hydrogen Bonds**

| Index | Residue | AA | Distance H-A | Distance D-A | Donor Angle | Protein donor? | Side chain | Donor Atom | Acceptor Atom |
|---|---|---|---|---|---|---|---|---|---|
| 1 | 26A | PHE | 3.02 | 3.45 | 106.68 | ✓ | × | 233 [Nam] | 10 [O3] |
| 2 | 390A | TYR | 2.27 | 3.14 | 147.98 | × | ✓ | 10 [O3] | 5856 [O3] |
| 3 | 390C | TYR | 2.09 | 3.00 | 155.77 | ✓ | ✓ | 13892 [O3] | 8 [O2] |
| 4 | 393A | GLN | 2.50 | 3.03 | 112.30 | ✓ | ✓ | 5909 [Nam] | 4 [O2] |
| 5 | 393C | GLN | 3.23 | 3.79 | 116.45 | ✓ | ✓ | 13945 [Nam] | 10 [O3] |
| 6 | 400C | ARG | 3.41 | 3.82 | 105.96 | ✓ | ✓ | 14076 [Ng+] | 2 [O2] |

**Figure S5: Shikonin inhibits nuclear translocation of PKM2 without altering its expression.** (A) Predicted interaction between PKM2 and shikonin. The structure of human PKM2 docked with shikonin. Inset shows the site pocket with surface charge distribution on donor and acceptor atoms. (B) The ligand-receptor interaction profile of PKM2-shikonin generated using PLIP. The aminoacid residues are shown in blue and shikonin is shown in orange. The hydrogen bond and hydrophobic interactions are represented by solid blue and dotted grey lines, respectively. PLIP output table demonstrating the expected interactions of PKM2 with shikonin *(top)* along with 3D visualization of the main interactions of the top predicted binding model *(bottom)* is provided. The $IC_{50}$ value of shikonin in WT BBS (C) MCF7 and (D) HCC1806 cells calculated under 24hrs of hypoxic exposure. (E) Immunoblot analysis of PKM2 expression upon shikonin treatment. For (E) representative image is provided. Error bars show mean values ± SD (n = 3 unless otherwise specified). As calculated using two-tailed Student's t-test, $^{*}P \leq 0.05$, $^{**}P \leq 0.01$, $^{***}P \leq 0.001$, $^{****}P \leq 0.0001$.

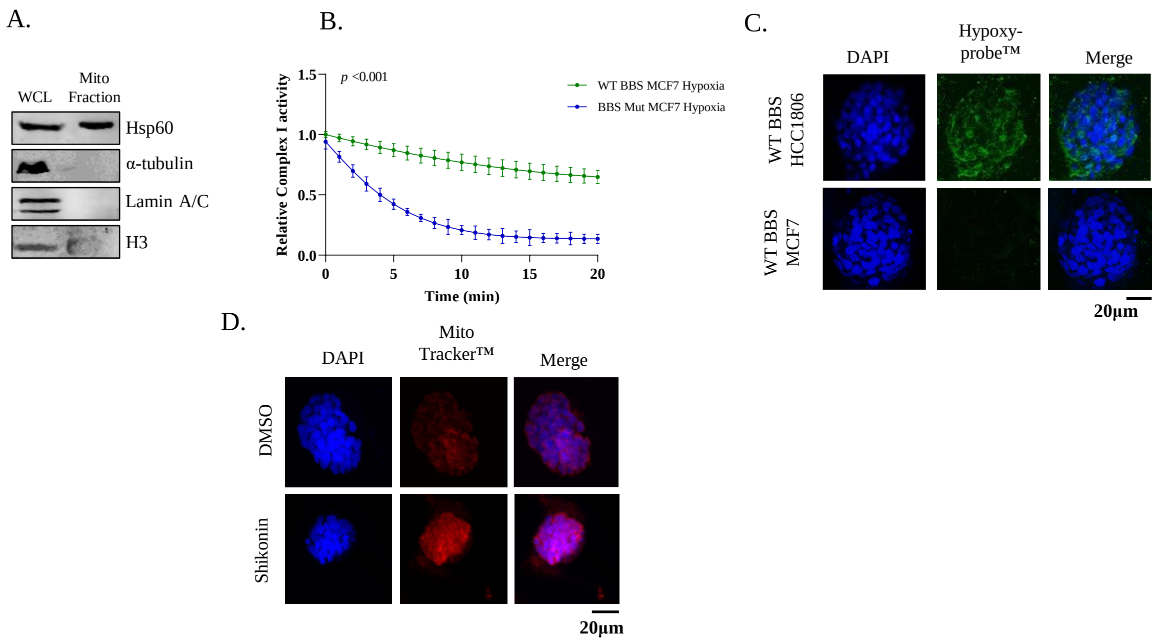


**Figure S6: Increased OXPHOS affects proliferation rate of breast cancer cells.** (A) Western blot showing the purity of mitochondrial fraction using α-tubulin (cytoplasmic marker), Lamin A/C and H3 (nuclear marker), and Hsp60 (mitochondrial marker) as controls. (B) Mitochondrial complex I activity assay. The assay indicates overall increased mitochondrial complex I activity in the BBS Mut MCF7 cells under hypoxia. (C) Hypoxyprobe™-1 immunofluorescence imaging performed in spheroids derived from WT BBS MCF7 and HCC1806. The Maximum intensity projection (MIP) images shows the presence of multiple hypoxic regions in the WT BBS HCC1806-derived-spheroids. Magnification used: 60X. (D) Fluorescence imaging performed in 3D spheroids derived from WT BBS HCC1806 cells showing enhanced MitoTracker™ Red uptake to indicate an increase in OXPHOS when treated with shikonin. Magnification used: 60X. Representative image are provided for all the figures. Error bars show mean values ± SD (n = 3 unless otherwise specified). As calculated using two-tailed Student's t-test, and unpaired t-test for mitochondrial complex I assay, $*P \leq 0.05$, $**P \leq 0.01$, $***P \leq 0.001$, $****P \leq 0.0001$.

### Table 1: Oligo Sequence of shRNAs and sgRNAs

| | |
|---|---|
| sh_Ctrl | 5’CCGGCGTGATCTTCACCGACAAGATCTCGAGATCTTGTCGGTGAAGATCACGTTTTT3’ |
| shHIF-1α_1 | 5’CCGGGTGATGAAAGAATTACCGAATCTCGAGATTCGGTAATTCTTTCATCACTTTTT3’ |
| shHIF-1α_2 | 5’CCGGTGCTCTTTGTGGTTGGATCTACTCGAGTAGATCCAACCACAAAGAGCATTTTT3’ |
| shHIF-2α_1 | 5’CCGGCGACCTGAAGATTGAAGTGATCTCGAGATCACTTCAATCTTCAGGTCGTTTTT3’ |
| shHIF-2α_2 | 5’CCGGCAGTACCCAGACGGATTTCAACTCGAGTTGAAATCCGTCTGGGTACTGTTTTT3’ |
| shRan_1 | 5’GTACCGGTGGACTGAGAGATGGCTATTACTCGAGTAATAGCCATCTCTCAGTCCATTTTTTG3’ |
| shRan_2 | 5’GTACCGGGAGTGAATGTGGCAGTTTAAACTCGAGTTTAAACTGCCACATTCACTCTTTTTTG3’ |
| shPKM2_1 | 5’CCGGGAACTCACTCTGGGCTGTAACCTCGAGGTTACAGCCCAGAGTGAGTTCTTTTTTG3’ |
| sgPKM2 | 5’TGGTAATGGGCGCCAGGCGG3’ |
| sgHIF-1α | 5′CTGTGATGAGGCTTACCATC3′ |
| sgHIF-2 α | 5’GATTGCCAGTCGCATGATGG3’ |

### Table 2: List of Primer Sequences

| | |
|---|---|
| PFKFB3_HRE1_Fw | 5’AGCTCGCAGGCTGCTTC3’ |
| PFKFB3_HRE1_Rv | 5’CGCAGACGCGTACGTCA3’ |
| PFKFB3_HRE2_Fw | 5’CCCTCCCTGTGGAGCAT3’ |
| PFKFB3_HRE2_Rv | 5’CGCTCGCTGAAACCTACAC3’ |
| PFKFB3_HRE3_Fw | 5’CTTGTAGGGAGTGTGGAAGAAG3’ |
| PFKFB3_HRE3_Rv | 5’TGCTTGCCCACCGATTT3’ |
| PFKFB1 Fw | 5’TAGGCCAGTATCGACGAG3’ |
| PFKFB1 Rv | 5’CCGTCGTTCTCTGGTAGT3’ |
| PFKFB2 Fw | 5’TGATCCTGATGTCATTGCTGC3’ |
| PFKFB2 Rv | 5’GTTGTCTGGGTCAAGAGG3’ |
| PFKFB3 Fw | 5’GTTAAAATCTCCAGCCCGGA3’ |
| PFKFB3 Rv | 5’AGGATCAGTTGCTATGAAGCC3’ |
| PFKFB4 Fw | 5’CAAACACCACCCGAGAAC3’ |
| PFKFB4 Rv | 5’CCGTAGCCTCATCACTGT3’ |
| SLC2A1 Fw | 5’TAAAGAAGCTGCGCGGGA3’ |
| SLC2A1 Rv | 5’CTTCTCGAAGATGCTCGTGGA3’ |

| | |
|---|---|
| SLC2A3 fw | 5’TCCCCTCCGCTGCTCACTATTT3’ |
| SLC2A3 Rv | 5’ATCTCCATGACGCCGTCCTTTC3’ |
| SLC16A3 Fw | 5’CCACAAGTTCTCCAGTGCCATTG3’ |
| SLC16A3 Rv | 5’CGCCAGGATGAACACGTACATG3’ |
| Aldo C Fw | 5’CATTCTGGCTGCGGATGAGTCT3’ |
| Aldo C Rv | 5’CACACGGTCATCAGCACTGAAC3’ |
| LDHA fw | 5’ATCTTGACCTACGTGGCTTGGA3’ |
| LDHA Rv | 5’CCATACAGGCACACTGGAATCTC3’ |
| HK1 Fw | 5’GGACTGGACCGTCTGAATGT3’ |
| HK1 Rv | 5’ACAGTTCCTTCACCGTCTGG3’ |
| HK 2 Fw | 5’CAAAGTGACAGTGGGTGTGG3’ |
| HK2 Rv | 5’GCCAGGTCCTTCACTGTCTC3’ |
| HRE1_Luc Fw | 5’TTTTGGTACCAGCTCGCAGGCTGCTTC3’ |
| HRE1-Luc Rv | 5’TTTTCTCGAGCGCAGACGCGTACGTCA3’ |
| HRE2_Luc Fw | 5’TTTTGGTACCGCGCCCTCCCTGTGGAGCAT3’ |
| HRE2_Luc Rv | 5’TTTTCTCGAGCGCTCGCTGAAACCTACAC3’ |
| HRE3_Luc Fw | 5’TTTTGGTACCCTTGTAGGGAGTGTGGAAGAAG3’ |
| HRE3_Luc Rv | 5’TTTTCTCGAGTGCTTGCCCACCGATTT3’ |
| PKM2 OE Fw | 5’TTTTGCGGCCGCATGTCACCGGAAGCCCAA3’ |
| PKM2 OE Rv | 5’TTTTGAATTCTCACGGCACAGGAACAAC3’ |
| RPS16 Fw | 5’AAACGCGGCAATGGTCTCATCAAG3’ |
| RPS16 Rv | 5’TGGAGATGGACTGACGGATAGCAT3’ |

### Table 3: List of antibodies

| S No. | Antibody | Company | Catalogue no. | Lot no. |
|---|---|---|---|---|
| 1 | PKM2 (D78A4) | Cell Signalling Technology | 4053S | 5 |
| 2 | PKM1 (D30g6) | Cell Signalling Technology | 7067S | 3 |
| 3 | HIF-1α (D2U3T) | Cell Signalling Technology | 14179S | 3 |
| 4 | HIF-2α (D6T8V) | Cell Signalling Technology | 59973S | 2 |
| 5 | Lamin A/C | Cell Signalling Technology | 2032S | 6 |
| 6 | GAPDH (D16h11) | Cell Signalling Technology | 5174S | 7 |
| 7 | Anti-PFKFB3 [EPR12594] | Abcam | ab181861 | GR3256208-2 |
| 8 | Anti-Ki67 | Abcam | ab197234 | GR205364-3 |

| | | | | |
|---|---|---|---|---|
| 9 | Hsp 60 | Cell Signalling Technology | 12165S | 4 |
| 10 | H3 | Active motif | 61475 | 25613001 |
| 11 | CAIX | Abcam | ab184006 | GR3388214-1 |
| 12 | α-Tubulin | Abcam | ab126165 | GR106527-20 |
| 13 | Alexa-Flour 680 anti-rabbit IgG | Invitrogen | A32734 | RJ243414 |
| 14 | Alexa-Flour 555 anti-rabbit IgG | Invitrogen | A32732 | 1858260 |
| 15 | Alexa-Flour 800 anti-mouse IgG | Invitrogen | A32730 | SC243837 |
| 16 | Alexa-Flour 488 anti-mouse IgG | Invitrogen | A11059 | 1832425 |
| 17 | Normal Rabbit IgG | Cell Signalling Technology | 2729S | 8 |
| 18 | HP1 (mAB for pimonidazole IgG clone 1) | Hypoxyprobe | 4.3.11.31 | 07.31.18 |
| 19 | P300 (D2x6n) | Cell Signalling Technology | 54062S | 1 |
| 20 | Mouse IgG isotype control | Invitrogen | 10400C | UG286589 |
| 21 | Histone H3 acetyl K9 | Abcam | ab10812 | GR3175803-1 |
| 22 | Tri-Methyl-Histone H3 (Lys9) (D4W1U) | Cell Signalling Technology | 13969S | 3 |

### Table 4: Clinical Characteristics of Patients

| S No. | Histopathology |
|---|---|
| 1 | Infiltrating duct carcinoma grade III Pathological stage pT2N1Mx |
| 2 | Infiltrating duct carcinoma Pathological stage ypT2N1Mx |
| 3 | Infiltrating duct carcinoma grade III Pathological stage pT2N0Mx |
| 4 | Infiltrating duct carcinoma grade III pathological stage pT2N1(mi)Mx |
| 5 | solid papillary invasive carcinoma left breast |
| 6 | Invasive carcinoma of no special type ductal grade II left breast pathological stage pT2N0Mx |
| 7 | Invasive carcinoma of no special type ductal grade II right breast pathological stage pT2N0Mx |
| 8 | Infiltrating duct carcinoma Grade II stage pT2N0Mx |
| 9 | Invasive carcinoma of no special type ductal with mucinous differentiation grade II right breast Pathological stage pT2N1Mx |
| 10 | Invasive carcinoma of no special type (ductal) grade II Pathological stage pT2NxMx |

| | |
|---|---|
| 11 | Invasive carcinoma of no special type (ductal) grade II of right breast Pathological stage pT2N1]Mx |
| 12 | Infiltrating duct carcinoma grade II of right breast Pathological stage pT4N1Mx |
| 13 | Infiltrating duct carcinoma grade III of right breast Pathological stage pT4N0Mx |
| 14 | Invasive carcinoma of no special type (ductal), grade II, Pathological stage pT2N0Mx |
| 15 | Invasive carcinoma of no special type (ductal), grade II, Pathological stage pT2N0Mx |
| 16 | Invasive carcinoma of no special type (ductal), grade III, Pathological stage pT2N3Mx |
| 17 | Invasive carcinoma of no special type (ductal), grade II, Pathological stage pT2N1Mx |
| 18 | Invasive carcinoma of no special type (ductal), grade I, Pathological stage pT1N0Mx |

**Table 5: Common genes in HIF-1α ChIP-seq and BBS Mut vs WT BBS MCF7 HTA 2.0 analyses**

| Genes | Fold Change |
|---|---|
| GDF15 | 18.22 |
| ATP9A | 11.19 |
| TFF1 | 7.76 |
| EGLN3 | 5.95 |
| MGAT5 | 4.65 |
| STC2 | 3.92 |
| IGFBP3 | 3.67 |
| PXDN | 3.64 |
| PPP1R3C | 3.55 |
| NCOA3 | 3.04 |
| TMEM45B | 3.03 |
| TRAM2 | 2.95 |
| NT5DC1 | 2.86 |
| PNPO | 2.79 |
| DHCR7 | 2.58 |
| RAI14 | 2.55 |
| ITGA5 | 2.51 |
| PTPRM | 2.42 |
| TRIM37 | 2.34 |
| HEY1 | 2.28 |
| C1orf116 | 2.26 |
| DNASE1 | 2.26 |
| TMEM45A | 2.17 |
| DSP | 2.16 |
| BZW1 | 2.13 |
| MAPK6 | 2.05 |
| NARF | 2.04 |

| | |
|---|---|
| SERTAD2 | -2.01 |
| LONP1 | -2.07 |
| PFKFB3 | -2.07 |
| WDR54 | -2.1 |
| PPP1CB | -2.15 |
| P4HA1 | -2.22 |
| PKP2 | -2.29 |
| WSB1 | -2.29 |
| SMC2 | -2.32 |
| ALDOC | -2.52 |
| KRT80 | -2.66 |
| CLK3 | -2.71 |
| ADM | -3.04 |
| CAV1 | -3.04 |
| FUT11 | -3.13 |
| HK2 | -4.29 |
| NDRG1 | -4.8 |
